# Choosing explanation over performance: Insights from machine learning-based prediction of human intelligence from brain connectivity

**DOI:** 10.1101/2023.12.04.569974

**Authors:** Jonas A. Thiele, Joshua Faskowitz, Olaf Sporns, Kirsten Hilger

**Affiliations:** Department of Psychology I - Biological Psychology, Clinical Psychology and Psychotherapy, Würzburg University, Marcusstr. 9-11, 97070 Würzburg, Germany; Department of Psychological and Brain Sciences, Indiana University, Psychology Building 360, 1101 E 10th Street, Bloomington, IN 47405, USA

**Keywords:** Machine Learning, Intelligence, Cognitive ability, Functional brain connectivity, Predictive modelling, Functional brain networks

## Abstract

A growing body of research predicts individual cognitive ability levels from brain characteristics including functional brain connectivity. The majority of this research achieves good prediction performance but provides limited insight into neurobiological processes underlying the predicted concepts. The insufficient identification of predictive characteristics may present an important factor critically contributing to this constraint. Here, we encourage to design predictive modelling studies with an emphasis on interpretability to enhance our conceptual understanding of human cognition. As an example, we investigated in a preregistered study which functional brain connections successfully predict general, crystallized, and fluid intelligence in a sample of 806 healthy adults (replication: *N* = 322). The choice of the predicted intelligence component as well as the task during which connectivity was measured proved crucial for better understanding intelligence at the neural level. Further, intelligence could be predicted not solely from one specific set of brain connections, but from various combinations of connections with system-wide locations. Such partially redundant, system-wide functional characteristics complement intelligence-relevant connectivity of brain regions proposed by established intelligence theories. In sum, our study showcases how future predictive studies on human cognition can enhance explanatory value by prioritizing a systematic evaluation of predictive characteristics over maximizing prediction performance.

**Significance Statement:** Intelligence represents a hallmark of human behavior, and a surge number of studies predicted individual scores from functional brain connectivity. However, actual understanding about its neural basis remains limited. We demonstrate how predictive modelling can be applied strategically to improve tracing predictive functional brain connections to enhance our understanding of intelligence. Our study unveils crucial findings about intelligence: differences in the neural code of distinct intelligence facets not detectable on a behavioral level and a brain-wide distribution of functional brain characteristics relevant to intelligence that extends those proposed by major intelligence theories. In a broader context, it offers a framework for future prediction studies that prioritize meaningful insights into the neural basis of complex human traits over predictive performance.

## Introduction

Neuroscientific research on human behavior and cognition has methodologically moved from unimodal explanatory approaches to machine learning-based predictive modelling (1). This implies a shift from standard approaches testing for associations between behavior and single neurobiological variables within one sample (unimodal explanatory research) to the identification of relationships between behavior and multiple neurobiological variables to forecast behavior of unseen individuals across samples (multimodal predictive research) (2). Modern machine learning techniques can learn such general relations in neural data (2, 3) and have consequently become increasingly prominent also in research on fundamental psychological constructs like intelligence (4).

Intelligence captures the general cognitive ability level of an individual person and predicts crucial life outcomes, such as academic achievement, health, and longevity (5, 6). Multiple psychometrical theories about the underlying conceptual structure of intelligence have been proposed. For example, Spearman (7) noticed that a person’s performance on different cognitive tasks is highly correlated and suggested that this ‘positive manifold’ results from an underlying common factor – general intelligence (*g*). A decomposition of the *g*-factor into fluid (*g*F) and crystallized (*g*C) components was later proposed by Cattell (8, 9). While fluid intelligence is assumed to mainly consist of inductive and deductive reasoning abilities that are rather independent of prior knowledge and cultural influences, crystallized intelligence reflects the ability to apply acquired knowledge and thus depends on experience and culture (10).

Neurobiological correlates of intelligence differences were identified in brain structure (11) and brain function (12). However, rather than disclose a single ‘intelligence brain region’, metanalyses and systematic reviews suggest the involvement of a distributed brain network (13–15), thus paving the way for proposals of whole-brain structural and functional connectivity underlying intelligence (16, 17). While the great majority of such studies used an explanatory approach, recently, an increasing number of machine learning-based techniques were developed and applied to predict intelligence from brain features (4, 18, 19). Although intrinsic functional connectivity measured during the (task-free) resting state has enabled robust prediction of intelligence (19), prediction performance can be boosted by measuring connectivity during task performance (18, 20).

However, despite extensive research, predictive modelling has provided only limited conceptual insights into human cognition and intelligence (4, 21). Reasons include the mostly restricted focus on the prediction of fluid intelligence, hindering examination of theories comprising multiple intelligence components, and the use of different tests to measure fluid intelligence (22, 23). Additionally, 32 % of studies reviewed in Vieira et al. (4) based their analyses on connectome-based predictive modeling (CPM) (24, 25). As CPM includes strict threshold-based feature selection, elimination of sub-threshold relevant information presents a concern (26). Last, previous studies strived for maximizing prediction performance, rather than for deriving predictive brain features exhaustively (4, 21). Although contributions of brain features were assessed, e.g., by estimating relative importance of model inputs by evaluating regression weights, such methods are criticized to lack reliability (21) and to provide misleading information (27, 28). Deeper insight into the relationship between functional brain connectivity and human intelligence that can also contribute to our conceptual understanding of human cognition therefore requires a systematic approach that compares predictions of different intelligence components, considers potential nonlinear relations, and includes a variety of different brain connectivity features selected on the basis of conceptual knowledge and theory.

Here, we aim to close this gap by providing exemplary means and methods to systematically examine the importance of brain characteristics (features) for predicting human traits. Specifically, we used functional connectivity (brain connections) of 806 healthy adults assessed during resting state and seven task states to predict general, crystallized, and fluid intelligence with non-linear machine learning models. We systematically estimated the contribution of different brain connectivity features a) by testing the predictive performance of single brain networks and network combinations with functional brain connection selection (19, 29), b) by comparing prediction performance from randomly selected functional brain connections with connections proposed as relevant by established intelligence theories, and c) by identifying a novel network of brain connections most critical for intelligence prediction using a modification of layerwise relevance propagation (LRP) – a method for estimating feature relevance (30). To ensure robustness and generalizability of findings, we cross-validated prediction models developed on the main part of the sample internally, in a lockbox sample, and in two independent samples.

## Results

### Intelligence and functional brain connectivity

General *g*, crystallized *g*C, and fluid *g*F intelligence components were estimated from 12 cognitive measures (*SI Appendix*, Table S1) for subjects of the Human Connectome Project (HCP) (31). Intelligence components were approximately normally distributed (*SI Appendix*, Fig. S1) and significantly positively correlated with each other: general and crystallized intelligence *r* = 0.78 (*p* < 0.001), general and fluid intelligence *r* = 0.76 (*p* < 0.001), crystallized and fluid intelligence *r* = 0.49 (*p* < 0.001). Individual functional connectivity (FC) between 100 cortical nodes (32) was constructed from resting state and from seven task states. Further, two latent FCs (33), one across rest and task states and one only across task states, were formed. This resulted in FCs of 10 conditions that are referred to as (cognitive) states. Group mean functional connectivity was highly similar across states and higher within than between different brain networks (*SI Appendix*, Fig. S2*A*). Between-subject variance in functional connectivity showed a network-specific pattern (*SI Appendix*, Fig. S2*B*). Descriptives of the combined samples (PIOP1, PIOP2) from the Amsterdam Open MRI collection (AOMIC) (34), which were used for replication, are illustrated in *SI Appendix*, Fig. S3.

### Functional connectivity better predicts general and crystallized intelligence than fluid intelligence

Performance to predict intelligence was investigated with a functional brain connection selection approach involving the systematic training and testing of prediction models with varying sets of brain connections (functional connections, Fig. 1) in the main sample (HCP, 610 subjects). Model performance was calculated as Pearson correlation (*r*) between predicted and observed intelligence scores and *p*-values were obtained with non-parametric permutation tests (uncorrected for the number of prediction models, FDR-corrected significances were mostly similar and are reported in *SI Appendix* Figs. S4-S10). First, all functional brain connections served as input features (whole-brain prediction). Averaged across all cognitive states, predictions of general, crystallized, and fluid intelligence reached statistical significance: prediction performance was highest for general intelligence (*r* = 0.31, *p* < 0.001), followed by crystallized intelligence (*r* = 0.27, *p* < 0.001), and fluid intelligence (*r* = 0.20, *p* < 0.001; Fig. 2). These differences were statistically significant: *g* vs. *g*C: *t*(9) = 2.77, *p* = 0.022; *g*C vs. *g*F: *t*(9) = 4.91, *p* < 0.001.

**Figure 1.**
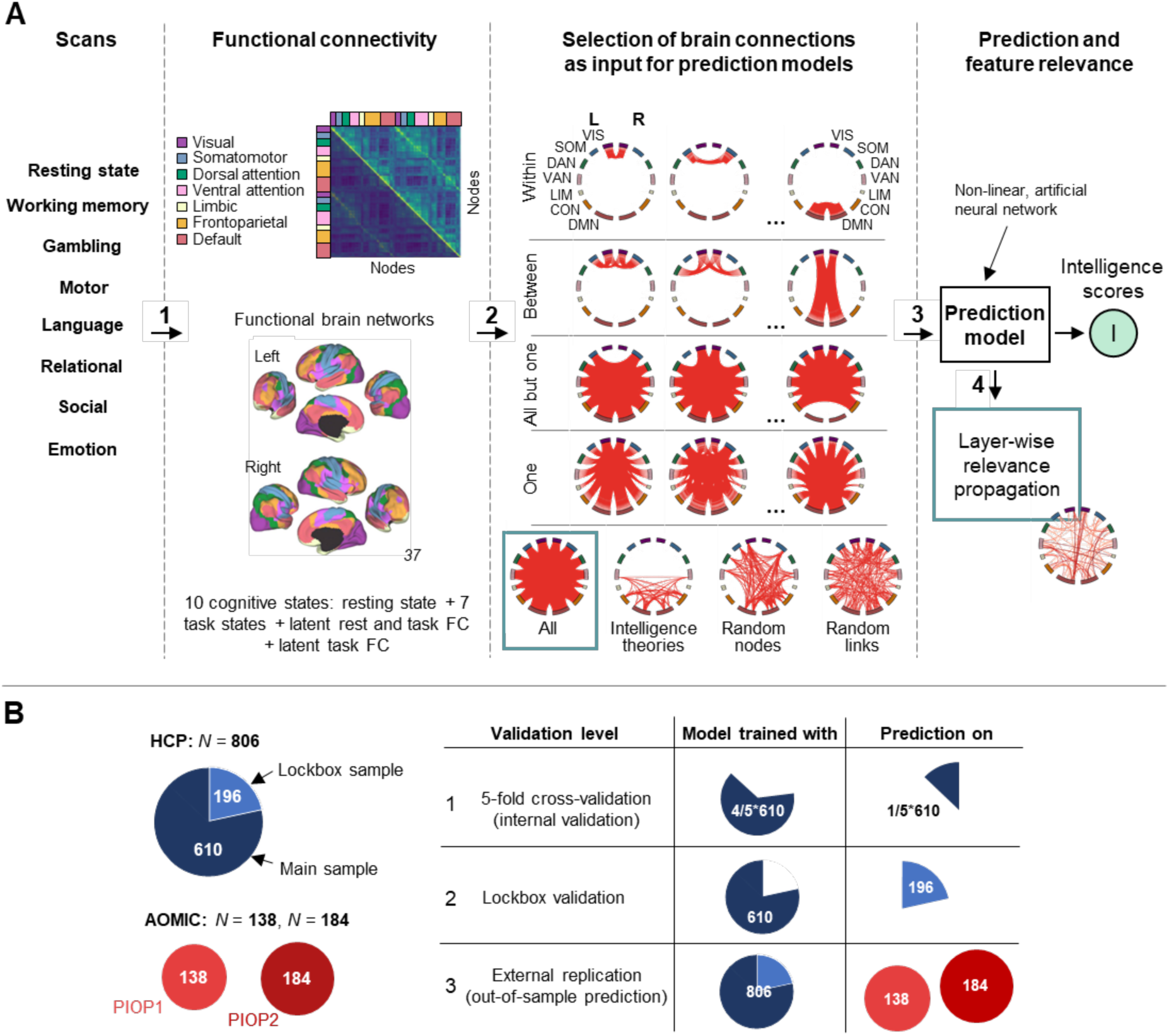
Schematic study overview. (A) Predicting intelligence scores from functional brain connectivity. 1 - Functional connectivity (FC) was estimated from fMRI data assessed during eight different cognitive states (resting state, seven tasks). Additionally, two latent FCs (33) were estimated based on a) resting state and all task states, and b) all task states. Functional brain connections were assigned to seven functional networks (35). 2 - Different selections of brain connections served as input features for prediction models, i.e., all connections, connections within a network or between two networks, connections of all but one network, all within- and between-network connections of one network, connections between brain regions (nodes) proposed as relevant by established intelligence theories, randomly selected brain connections, and connections between randomly selected nodes. 3 - Prediction of general *g*, crystallized *g*C, or fluid *g*F intelligence with models, separately trained with the different selections of brain connections. 4 – Estimation of connection-wise contributions to the prediction of intelligence in models trained with all brain connections via stepwise layer-wise relevance propagation. (B) Overview of study samples and cross-validation. The Human Connectome Project (HCP) sample was first divided into a main (610 subjects) and a lockbox (196 subjects) sample. Main analyses were conducted with 5-fold cross-validation (validation step 1, internal validation) in which models are trained on four subsamples of the main sample and tested on the withheld fifth subsample. Second, models were trained on the main sample and used to predict intelligence scores in the lockbox sample (validation step 2, lockbox validation). Lastly, models trained on the HCP sample were used to predict intelligence scores in two independent samples of the Amsterdam Open MRI Collection (AOMIC: PIOP1 and PIOP2, validation step 3, external replication).

**Figure 2.**
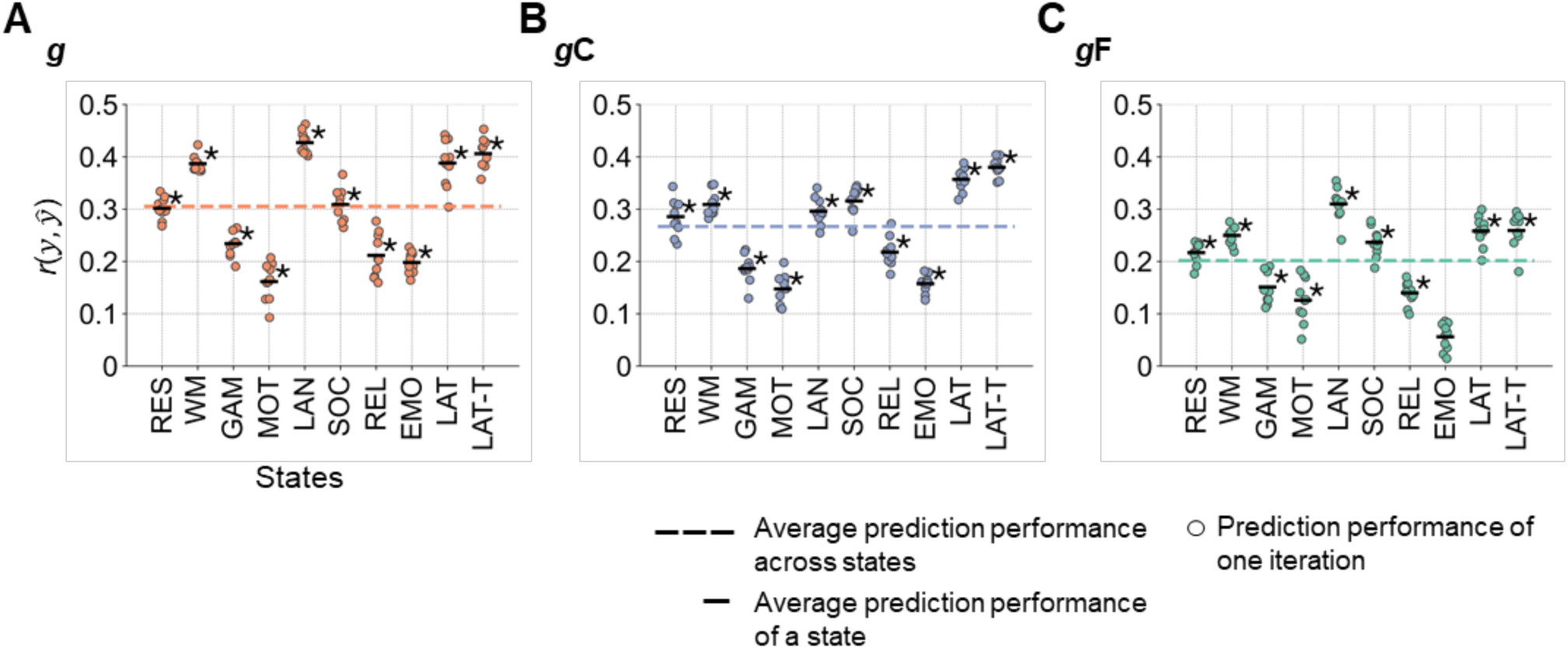
Performance of predicting intelligence from whole-brain functional connectivity. (A) Prediction of general *g*, (B) crystallized *g*C, and (C) fluid *g*F intelligence in the main sample (*N*=610, HCP). Prediction performance was calculated as Pearson correlation between observed and predicted intelligence scores *r*(*y*, *y*&). The performance of 10 prediction models trained with varying stratified folds is illustrated with colored dots (general intelligence red, crystallized intelligence blue, fluid intelligence green). The mean performance across these 10 models is indicated by the black horizontal bar. The mean performance across states is highlighted with a colored dashed line. Significant mean prediction performance (*p* < 0.05, permutation test, 100 permutations) is marked with an asterisk (uncorrected for the number of prediction models; FDR-corrected significances are reported in *SI Appendix* Fig. S4). RES, resting state; WM, working memory task; GAM, gambling task; MOT, motor task; LAN, language processing task; SOC, social cognition task; REL, relational processing task; EMO, emotion processing task; LAT, latent functional connectivity of resting state and all task states; LAT-T, latent functional connectivity of all task states.

### Prediction performance depends on cognitive brain states

All cognitive states except the emotion task allowed for significant predictions of all three intelligence components (Fig. 2). Error measures (*SI Appendix*, Fig. S11, see Methods) indicated comparable patterns. General intelligence was significantly better predicted from functional connectivity of the language task, the working memory task, and both latent functional connectivity factors than by the gambling, the relational, the emotion, and the motor task (model difference test: all *p* < 0.05). Crystallized intelligence was best predicted from both latent functional connectivity factors, followed by the social and working memory task, while predictions from the emotion and motor task were significantly worse (model difference test: all *p* < 0.05). Most predictive states for fluid intelligence were the language task, followed by latent functional connectivity factors, the working memory and the social task all of which performed significantly better than the emotion task (model difference test: *p* < 0.05).

### Brain networks differ in their ability to predict intelligence

Next, the ability to predict intelligence from different functional brain networks (35) as well as from different network combinations was investigated. Three selections of brain connections served as input features: a) all functional brain connections within a specific network or between two specific networks, b) all connections (whole-brain) but those of one specific network, and c) all within- and between-network connections of one specific brain network.

Across selections (a,b,c), general intelligence was predicted significantly best, while fluid intelligence was predicted significantly worst (all *p* < 0.05, paired *t*-test between components for each selection approach; Figs. 3,4; for error measures: *SI Appendix*, Figs. S12-S15). Further, connectivity of the language and the working memory task, as well as both latent connectivity factors significantly outperformed connectivity of the relational, emotion, and gambling task (*p* < 0.05, significant differences between the prediction performance of two states, paired *t*-tests for each selection approach). Both confirmed the whole-brain prediction results.

**Figure 3.**
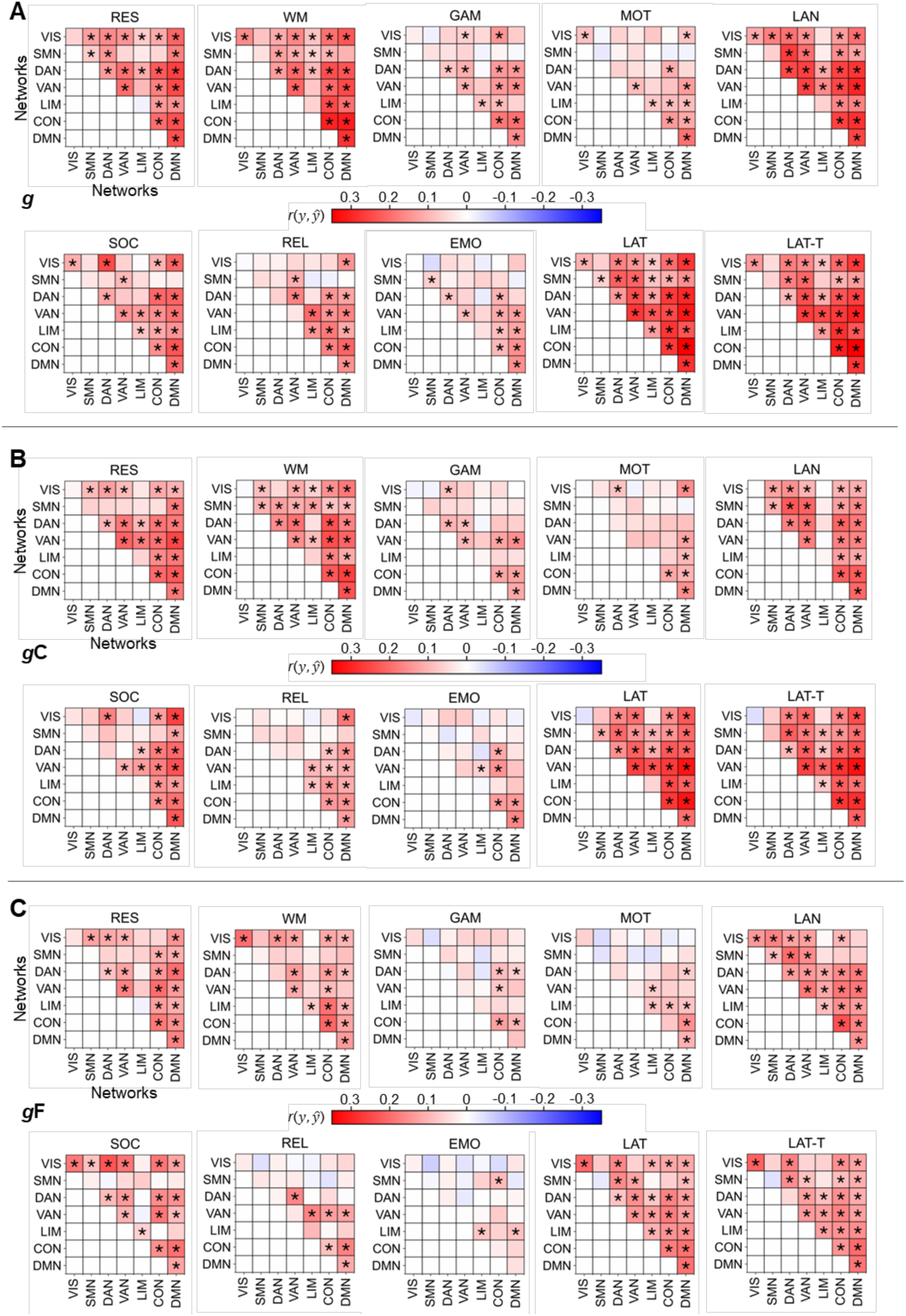
State- and network-specific performance of intelligence prediction from functional brain connections within a specific network or between two specific networks. (A) Prediction of general g, (B) crystallized gC, and (C) fluid gF intelligence in the main sample (N=610, HCP). Prediction performance (Pearson correlation between observed and predicted intelligence scores *r*(*y*, *y*&)) was calculated as average across 10 models trained with varying stratified folds. Significant prediction performance (*p* < 0.05, permutation test, 100 permutations) is marked with an asterisk (uncorrected for the number of prediction models; FDR-corrected significances are reported in *SI Appendix* Fig. S5). RES, resting state; WM, working memory task; GAM, gambling task; MOT, motor task; LAN, language processing task; SOC, social cognition task; REL, relational processing task; EMO, emotion processing task; LAT, latent functional connectivity of resting state and all task states; LAT-T, latent functional connectivity of all task states; VIS, visual network; SMN, somatomotor network; DAN, dorsal attention network; VAN, salience/ventral attention network; LIM, limbic network; CON, control network; DMN, default mode network.

**Figure 4.**
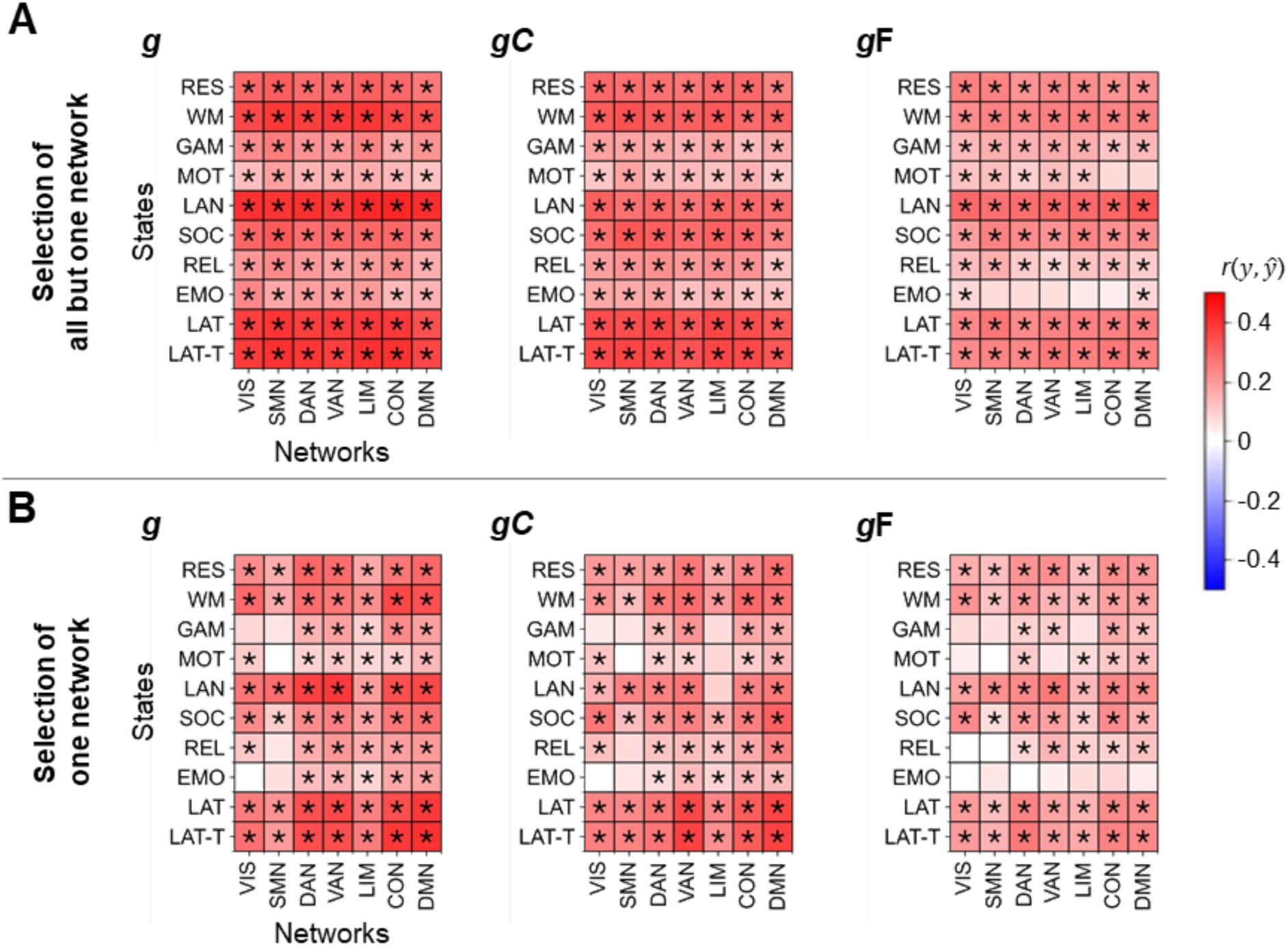
Performance of predicting intelligence from all functional brain connections but those of one specific brain network versus from connections of one brain network only. Models for predicting general *g* (left panels), crystallized *g*C (center panels), or fluid *g*F (right panels) intelligence in the main sample (*N*=610, HCP). (A) Separate models were trained with all connections but those of one specific network. (B) Models were trained with connections of one specific network. Prediction performance (Pearson correlation between observed and predicted intelligence scores *r*(*y*, *y*&)) was calculated as average across 10 models trained with varying stratified folds. Significant prediction performance (*p* < 0.05, permutation test, 100 permutations) is marked with an asterisk (uncorrected for the number of prediction models; FDR-corrected significances are reported in *SI Appendix* Fig. S6). RES, resting state; WM, working memory task; GAM, gambling task; MOT, motor task; LAN, language processing task; SOC, social cognition task; REL, relational processing task; EMO, emotion processing task; LAT, latent functional connectivity of resting state and all task states; LAT-T, latent functional connectivity of all task states; VIS, visual network; SMN, somatomotor network; DAN, dorsal attention network; VAN, salience/ventral attention network; LIM, limbic network; CON, control network; DMN, default mode network.

Focusing on the level of brain networks, selection (a) revealed that despite overall better prediction of general and crystalized intelligence compared to fluid intelligence, the patterns of which networks or network combinations perform better and which worse were relatively similar across intelligence components (Pearson correlation between the performance of all within- and between-network combinations predicting *g* vs. *g*C: *r* = 0.87, *p* < 0.001; *g* vs. *g*F: *r* = 0.79, *p* < 0.001; and *g*F vs. *g*C: *r* = 0.61, *p* < 0.001). Averaged across states, highest prediction performance was achieved from the default, control, and both attention networks, while visual, somatomotor, and limbic networks performed worse (mean performance over all within- and between-network combinations a respective network was involved in, averaged across all states). For the prediction of general and crystallized intelligence this effect reached statistical significance (paired *t*-tests for each combination of two networks, all *p* < 0.05). Similar findings were observed for predictions from all within- and between-network connections of one specific network (selection c) with the default, control, and both attention networks outperforming somatomotor and limbic networks (significant for general, crystallized, and fluid intelligence: paired *t*-tests for each combination of two networks, all *p* < 0.05). Note however, that despite these general trends some specific network combinations differed markedly in their predictive performance (*SI Appendix*, Fig. S16).

Finally, selections (b) and (c) revealed a high ability for compensating intelligent-relevant information: for all intelligence components, prediction performance did not significantly deviate from whole-brain prediction when excluding one brain network (selection b). For most cognitive states this even holds true when excluding all brain connections but those of only one brain network (selection c). Note that the latter refers only to ‘cognitive brain networks’ i.e., the default, control, or an attention network (model difference tests between models with all connections and models with all but one network or models trained with one network, Fig. 4, for error measures: *SI Appendix*, Fig. S15).

To rule out that results depend on the selected brain parcellation, we repeated the prediction of general intelligence from whole-brain and network-specific connectivity using the 200 node parcellation of Schaefer et al. (32). Using the finer parcellation, the prediction performance averaged across all states and that of selections a-c was slightly but significantly higher than the performance of the 100 node parcellation (200 nodes: *r* = 0.18; 100 nodes: *r* = 0.17; *p* < 0.05). Also, the pattern of the prediction performance across different brain networks, network combinations and cognitive states was highly similar (Pearson correlation between the prediction performance of all 10 (states) x 43 (connection selections) = 430 models: *r* = 0.94, *p* < 0.001). To evaluate the influence of socioeconomic status, we predicted general intelligence from whole-brain and network-specific connectivity with household income (HCP variable: SSAGA_Income) as additional control variable. Again, the pattern of prediction performance was highly similar (*SI Appendix*, Fig. S17, Pearson correlation between the prediction performance of all 10 (states) x 43 (connection selections) = 430 models: *r* = 0.97, *p* < 0.001). Lastly, to rule out that differences in prediction performance of different brain networks result merely from differences in the extent to which these networks capture individual differences, i.e., generally vary between persons (36), we tested for potential influences of inter-individual variances in connectivity strength of network-specific brain connection selections on their prediction performance. Although a positive trend emerged, it did not reach statistical significance for most states (*SI Appendix*, Fig. S18).

In sum, different approaches of functional brain connection selection revealed that the performance to predict intelligence depends on intelligence components (general intelligence best, fluid intelligence worst), cognitive states (cognitively demanding states and latent connectivity best), and brain networks (cognitive networks outperform other networks), while all brain networks contain intelligence relevant information and predictions are only minimally affected by removing entire brain networks.

### Support for neurocognitive models of intelligence

To inform established intelligence theories, we implemented a fourth selection approach and predicted general intelligence from functional brain connections between brain regions (node clusters) proposed by the revised Parieto-Frontal Integration Theory (P-FIT) (13), the multiple demand theory (MD) (14, 37), and the lateral PFC hypothesis (LPFC) (38). The resulting prediction performance was tested against null models, which were trained on an equal number of brain connections between an equal number of nodes, while nodes were selected randomly (100 permutations). This ensured that the null models had the same number of features, but with nodes distributed across the entire cortex. Although theory-driven models predicted worse than whole-brain models (Figs. 2*A*, 5*F*), they predicted significantly better than null models in multiple cases, while performance generally improved with increasing numbers of connections (Fig. 5). These results support a brain-wide distribution of intelligence-predictive information and show that theoretically proposed brain regions, although not being the sole determinants, significantly contribute to the prediction of intelligence.

**Figure 5.**
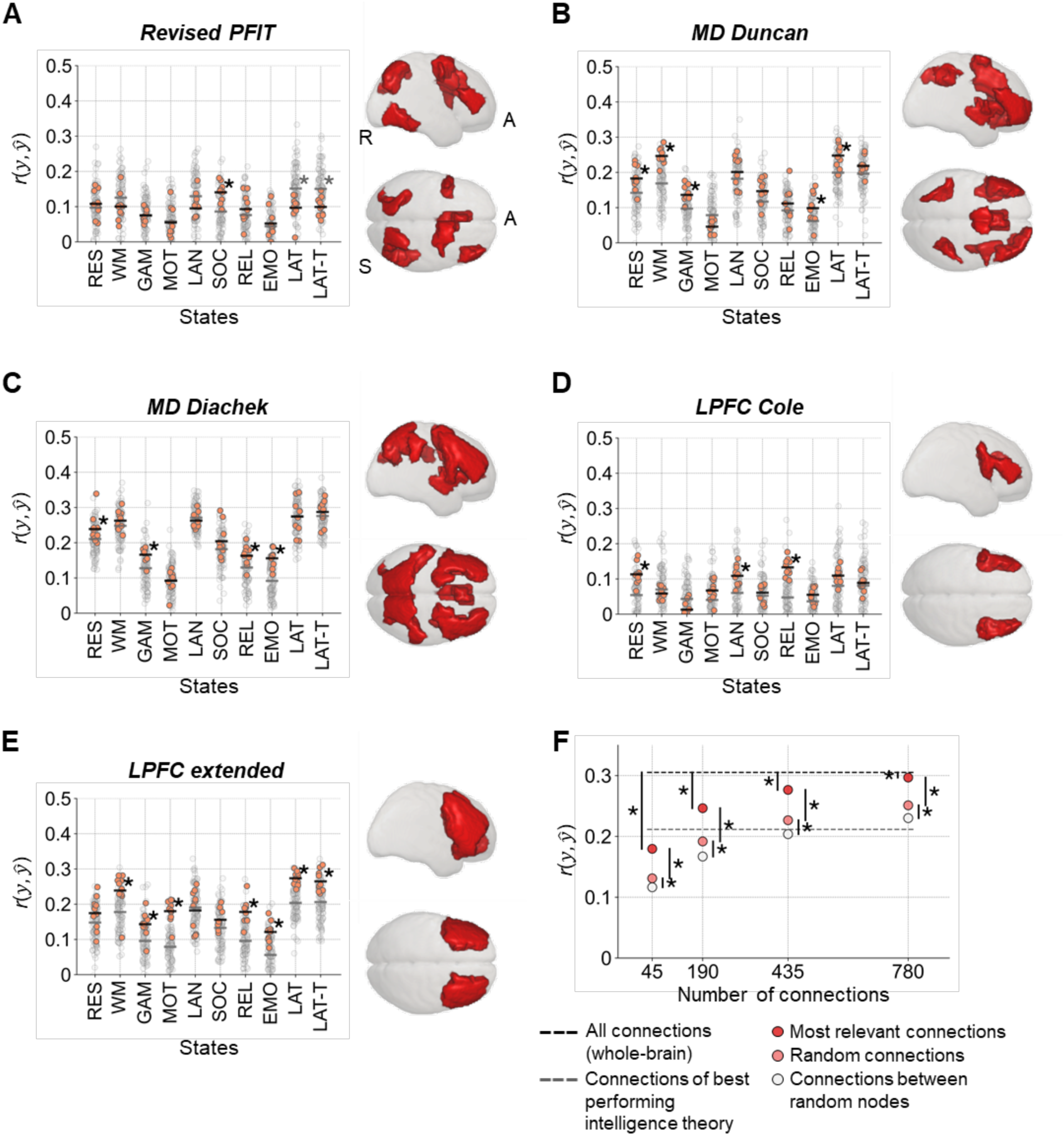
Performance of predicting general intelligence from theory-driven selections of functional brain connections. (A-E) The performance of models trained with connections between brain regions (nodes) proposed by established intelligence theories was tested against the performance of models, which were trained on an equal number of functional brain connections between an equal number of randomly selected nodes (null models). The performance of single theory-driven models (varying stratified folds) is displayed by a red dot, the mean performance across those models by the black horizontal bar. The performance of single null models is illustrated with a gray dot, their mean performance with a grey horizontal bar. Significant prediction performance (*p* < 0.05, permutation test, 100 permutations) is marked with an asterisk. Models were trained with all brain connections between (A) eight nodes of the revised P-FIT (13), (B) 13 nodes of the MD system (14), (C) 26 nodes of the MD system (37), (D) four LPFC nodes (38), and (E) an extension of LPFC nodes (14 nodes). Node cluster locations for each theory are illustrated on the right. (F) State-average prediction performance of models trained with different numbers of most relevant brain connections (red circles), randomly selected connections (light red circles), and connections between randomly selected nodes (white circles). The black dashed line reflects state-average performance of models trained with all brain connections. The gray dashed line illustrates state-average performance of models trained with connections from the best performing intelligence theory, MD by Diachek et al. (37) (for clarity, significant differences to other model performance is not displayed). Prediction performance was calculated as Pearson correlation between observed and predicted intelligence scores *r*(*y*, *y*&). Significant differences (*p* < 0.05, paired *t*-test) are marked with asterisks (uncorrected for the number of prediction models). RES, resting state; WM, working memory task; GAM, gambling task; MOT, motor task; LAN, language processing task; SOC, social cognition task; REL, relational processing task; EMO, emotion processing task; LAT, latent functional connectivity of resting state and all task states; LAT-T, latent functional connectivity of all task states; A, anterior; S, superior; R, right.

Given the brain-wide distribution of intelligence-relevant information, and the dependence of prediction performance on the number of brain connections, we next implemented a fifth selection approach and compared the prediction performance of models trained with different numbers of randomly selected connections with models trained with all possible connections between randomly selected nodes. Models based on randomly selected brain connections outperformed models based on connections between randomly selected nodes (Fig. 5*F*), while prediction performance generally increased with increasing numbers (45 to 780) of brain connections, approaching but not reaching the performance of whole-brain prediction. This underscores the assumption of a large, distributed network of intelligence-relevant brain connections and raises the question which and how many brain connections are required to enable the best possible prediction performance.

### Intelligence is best predicted from a widely distributed connectivity network

To identify a network of functional brain connections best predicting intelligence, we estimated connection-specific contributions using stepwise layer-wise relevance propagation (LRP). This allows for ascertaining contributions of single input features to predictions based on backpropagation (see Methods), here the identification of most relevant brain connections. The performance of prediction models trained with different numbers of these most relevant connections was then compared. Overall, models trained with most relevant connections outperformed models trained with randomly selected connections (Fig. 5*F*), while whole-brain performance was reached at 1000 connections for general and crystallized but not for fluid intelligence (no significant differences between the performance of models based on most relevant connections vs. all connections, Fig. 6*A*).

**Figure 6.**
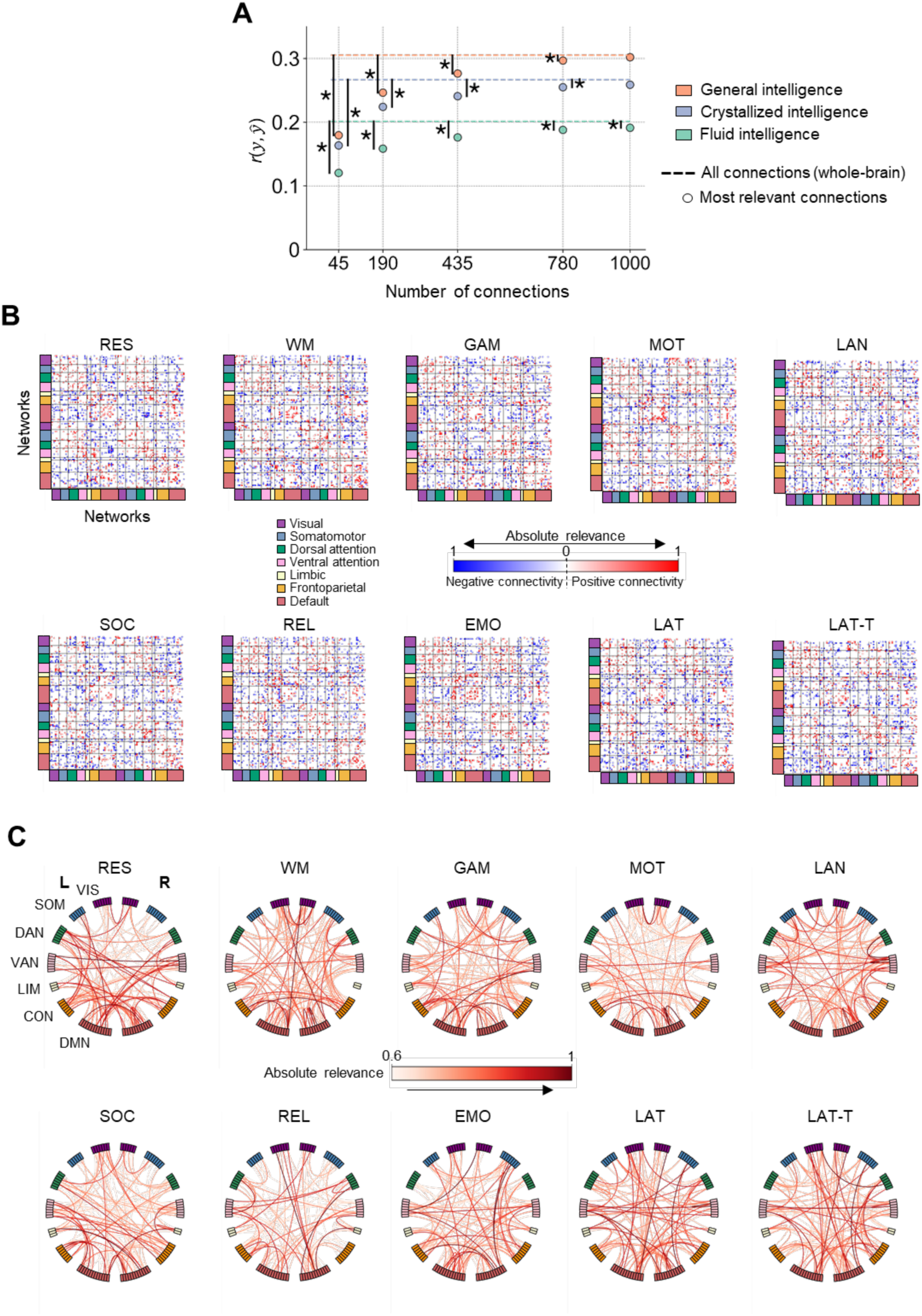
Intelligence is best predicted from a data-driven selection of 1000 most relevant functional brain connections defining a widely distributed network. (A) Performance of predicting general (red dots), crystallized (blue dots), and fluid (green dots) intelligence from models trained with different numbers of most relevant connections identified by stepwise layer-wise relevance propagation (LRP; N=610, HCP). Prediction performance was calculated as average of Pearson correlations between observed and predicted intelligence scores *r*(*y*, *ŷ*) across all states and 10 iterations (varying stratified folds). Dashed lines indicate state-average prediction performance of models trained with all functional connections (Fig. 2). Significant differences (*p* < 0.05, paired t-test) between the prediction performance of specific numbers of most relevant connections and whole-brain predictions are marked with asterisks. (B) Matrices of the 1000 most relevant connections. Red and blue indicate whether the mean strength of a functional brain connection is positive or negative (across subjects), while the saturation of the color displays a connection’s relevance (averaged over 10 iterations with varying stratified folds). (C) Connectograms of most relevant connections (for clarity, only the 100 most relevant brain connections are displayed; averaged over 10 iterations with varying stratified folds). RES, resting state; WM, working memory task; GAM, gambling task; MOT, motor task; LAN, language processing task; SOC, social cognition task; REL, relational processing task; EMO, emotion processing task; LAT, latent functional connectivity of resting state and all task states; LAT-T, latent functional connectivity of all task states; VIS, visual network; SMN, somatomotor network; DAN, dorsal attention network; VAN, salience/ventral attention network; LIM, limbic network; CON, control/frontoparietal network; DMN, default mode network.

The 1000 most relevant brain connections were widely distributed across the entire cortex and varied markedly between states with 19-27 % overlap between tasks, 28-38 % between tasks and latent connectivity, and 71-75 % between both latent connectivity factors (within each intelligence component; Figs. 6*B*,*C*, *SI Appendix*, Figs. S19, S20). The overlaps in the 1000 most relevant connections between the three intelligence components (across states) were 45 % between *g* and *gC*, 49 % between *g* and *g*F, and 28 % between *g*C and *g*F. Post-hoc analyses revealed that the 1000 most relevant connections did not systematically differ from randomly selected brain connections (1000 permutations) neither in their correlations with age, sex, handedness, or head motion nor in their test-retest reliability (ICC between FC of RL and LR phase) (39). In contrast, nodes connected to the 1000 most relevant connections had lower within-module degree z-scores and a tendency of higher participation coefficients (average of products of the number of occurrences of each node and the corresponding module-degree z-score or participation coefficient across all subjects) compared to randomly selected connections (1000 permutations, *p* < 0.05). This was observed across all states.

### Lockbox validation and external replication

For lockbox validation, we applied prediction models trained on the main sample (*N*=610, HCP) to a withheld subsample of the HCP (*N*=196). The prediction performance of the lockbox sample was significantly correlated with that of the main sample (Pearson correlation between the performance of whole-brain and network-specific predictions). This holds for general (*r* = 0.89, *p* < 0.001), crystallized (*r* = 0.84, *p* < 0.001) and fluid (*r* = 0.70, *p* < 0.001) intelligence (*SI Appendix*, Figs. S21-S23). The 1000 most relevant brain connections identified in the main sample predicted intelligence also significantly better in the lockbox sample than random connections, and connections between randomly selected nodes performed significantly worst. Again, prediction performance increased with increasing numbers of brain connections (*SI Appendix*, Fig. S24).

For external replication, we used models trained on the main sample (HCP) to predict intelligence scores in two combined samples of the AOMIC. Due to different fMRI tasks and intelligence assessments, only HCP’s *g* and *g*F models from: a) resting-state functional connectivity, b) task-connectivity (working memory task, emotion task), c) latent connectivity of resting state and all task states, and d) latent connectivity of all task states were tested for transferability. Prediction performance was substantially lower in the replication sample, but results between both samples (performance of whole-brain and network-specific predictions) were significantly correlated (*g*: *r* = 0.73, *p* < 0.001; *g*F: *r* = 0.66, *p* < 0.001; *SI Appendix*, Fig. S25). The 1000 most relevant brain connections identified in the HCP predicted intelligence also in the replication sample significantly better than random connections, while connections between random nodes, again, performed significantly worst, and prediction performance increased with increasing numbers of connections (*SI Appendix*, Fig. S24).

In sum, the lockbox validation proposes robustness of our results, while the result of the external replication demonstrates that, although effect sizes are lower, the overall patterns of results can be reproduced – despite differences in data acquisition parameters, preprocessing pipelines, and intelligence assessments in the replication sample.

## Discussion

Our study builds on extensive research predicting complex human traits from neuroimaging data. At first, we replicated that general, crystallized, and fluid intelligence can be predicted from functional brain connectivity measured during different cognitive states. Then, we demonstrated that predictions were significantly better for general and crystallized intelligence than for fluid intelligence, although only the latter was mostly addressed in previous work. Further, prediction results varied systematically between different states and brain networks associated with cognitive functions predicted best. Finally, different approaches of selecting functional brain connections revealed that brain-wide connectivity predicted significantly best, connectivity between brain regions proposed by leading intelligence theories predicted in multiple cases better than an equal number of connections between randomly chosen nodes, and that the impact of excluding complete functional networks was remarkably low.

Across all analyses, prediction performance varied systematically between intelligence components. This is remarkable, as individual contributions of cognitive states and functional brain networks were relatively similar between general, crystallized, and fluid intelligence and all three components were highly correlated on the behavioral level (*r* = 0.76 - 0.79) (40, 41). The observed differences in prediction performance suggest distinctions in components’ neural underpinnings which elude detection on a purely behavioral level (42) and inform future work to systematically evaluate their shared and specific neural correlates, also in comparison to existing intelligence theories (e.g., the process overlap theory) (43). In this context, it might be instructive to consider more delineated cognitive processes (41, 44) like memory capacity (45), attentional control (46), and processing speed (47, 48) to enlighten their involvement in general intelligence from a neural perspective.

General intelligence was predicted best, followed by crystallized intelligence and fluid intelligence. This pattern aligns with the systematic review of Vieira et al. (4), who observed a consistent trend of better prediction of general compared to fluid intelligence, however, it contrasts their challenging distinction at the behavioral level (40, 41). A potential reason for the lower predictability of fluid intelligence is the lower validity of fluid intelligence’s measurements, especially when measured by single tests (4, 49). We attempted to estimate fluid intelligence more validly with a composite score of different cognitive tasks but observed the same effect. Besides low-quality measurement of (fluid) intelligence, the observed pattern could also result from differing neural correlates like those that have been found in brain structure and function, specifically between fluid and crystallized intelligence (50–52). However, variations in the underlying neural processes remain largely unknown. We speculate that neural strategies, which involve distinct sets of functional brain connections, underlying crystallized and general intelligence may be more similar between individuals than those of fluid intelligence, rendering general and crystallized intelligence more predictable. For example, crystallized intelligence may be based on general knowledge stored in brain-wide fragments (knowledge networks) (51) with relatively similar localization and retrieval strategies. In contrast, fluid intelligence may involve complex interactions of different processes, such as working memory, attention, visuo-spatial reasoning (50) and processing speed (48), each of which may vary more strongly between individuals than the neural implementation of knowledge networks. The present study cannot differentiate between both explanations (low-quality measurement vs. more variable neural substrates), however, this question is an interesting subject for future research.

General and fluid intelligence were best predicted from functional connectivity assessed during the language task, while crystallized intelligence was best predicted by latent connectivity calculated from all states. Task-induced connectivity outperforming resting-state connectivity has also been observed in previous intelligence prediction studies (18), particularly when tasks induce high cognitive load (20). Our study additionally suggests that this dependence on cognitive load refers specifically to the prediction of fluid and general intelligence but less to crystallized intelligence. Speculatively, tasks requiring higher cognitive effort induce changes in functional connectivity that support inductive and deductive reasoning thereby enhancing the prediction of fluid intelligence. In contrast, crystallized intelligence might be best reflected in task-general connectivity characteristics represented in latent functional connectivity (33). Thus, a hypothesis that requires future investigation would be that crystallized intelligence is primarily coded in latent communication patterns possibly reflecting static structural brain characteristics that form a brain-wide knowledge network (51), whereas fluid intelligence relies on more specific neural communication processes that become particular visible during cognitively demanding tasks. If this holds true, it would indicate a differentiability between processes of crystallized and fluid intelligence at the level of macroscale hemodynamics, which can be made observable by adequate task selection.

Addressing the insufficient identification and systematic investigation of predictive brain characteristics in previous work, we developed an approach to predict intelligence from different selections of functional brain connections. Specifically, we show that significant prediction of intelligence was possible alone with connectivity of most individual networks as well as with connectivity between most combinations of two specific networks. The ability to compensate for missing intelligent-relevant connections was, thus, remarkably high and suggests that similarly successful prediction models can be constructed from different combinations of input features. Notably, this hinders the determination of *all* intelligence-related connections post-hoc with relevance estimation methods such as feature weight interpretation (21, 27, 28). Means like systematic functional brain connection selection can partially overcome this problem.

Despite brain-wide distribution of intelligence-relevant functional connections, the default mode, the fronto-parietal control, and both attention networks showed highest predictive power, while the somatomotor and limbic networks were least predictive. This coincides with previous research. First, functional connectivity *within* the fronto-parietal network for cognitive control and executive functioning has widely been associated with fluid intelligence (53) and general cognitive performance (19, 20) and also the default mode network was stressed in previous studies (19). The importance of connectivity within attentional brain systems for intelligence has been demonstrated in previous work (54–56), proposing mechanisms of attentional control, salience processing and the filtering of irrelevant information crucial for intelligence differences. Second, also connectivity *between* different networks was suggested to play an important role in cognitive processing. Particularly, the anti-correlation between task-positive (frontoparietal and attention networks) and task-negative (default mode) networks has been related to intelligence (20, 57, 58). Our results confirm the predictive potential of this between-networks interplay and undermine the assumption that counter-regulation of task-relevant and task-irrelevant processes is essential for cognitive functions relevant to intelligence. Based on these considerations, we recommend future research to disentangle the importance of fronto-parietal, default-mode, and attention networks for intelligence more in detail a) by testing differences in networks’ abilities to predict more circumscribed intelligence-related abilities involving processes like working memory, attentional control and processing speed, b) by designing in-scanner tasks in a way that they differ in their demands on such specific cognitive abilities (e.g., tasks requiring different degrees of attention), and c) by exploring differences in network’s relevance between healthy individuals and patients with impairments in specific cognitive functions (e.g., hemineglect).

In applying theory-driven selections of functional brain connections, we demonstrated that models trained with connections between brain regions proposed in neurocognitive intelligence theories (P-FIT (13, 15), MD (14, 37), LPFC (38)) outperformed models trained with the same number of connections between randomly chosen regions in several cases. Particularly, theories including an extended area around the lateral prefrontal cortex seemed to excel in prediction performance, thus, serving empirical support for these theories. Together with the observation that theory-driven selections performed significantly worse than models trained with all brain connections (59), our results suggest that connections between theoretically proposed brain regions include important but not complete intelligence-predictive information.

The identification of brain connections most relevant for intelligence prediction by stepwise LRP revealed that approximately 1000 connections are required to achieve a similar performance as whole-brain models. Those most relevant brain connections differed between cognitive states and varied for fluid, crystallized, and general intelligence. However, regardless of state and intelligence component, most relevant functional brain connections span a distributed network involving all 100 analyzed nodes and all major functional brain systems. Notably, the 1000 most relevant connections significantly outperformed randomly selected connections, confirming that indeed some connections are more relevant to intelligence prediction than others. Nodes connected to these connections were characterized by higher participation coefficients thus facilitating information transfer between brain modules and lower within-module degree z-scores reflecting less hub-like character (60).

Together, the good performance of randomly selected connections and the system’s high compensatory ability suggest the existence of a certain degree of redundancy in intelligence relevant information within functional brain connectivity (19). A potential cause for such redundancy may be that important traits like intelligence involve acting over different neural pathways possibly reflecting diverse cognitive strategies. This hypothesis aligns with studies indicating that neural redundancy plays a protective role in cognitive aging (61) and neurodegeneration (62), and enhances neural computation (63). Relatedly, a greater brain ‘resilience’ was observed in people with higher cognitive ability (64, 65).

Our study has several limitations. First, we only included three broad intelligence components (*g, g*C*, g*F). The consideration of additional or more circumscribed sub-components of intelligence may provide supplementary insights into its neural bases, specifically, into involved behavioral and neural subprocesses and strategies. Second, our analyses were restricted to static functional connectivity. However, the dynamics of neural processes could also include intelligence-predictive information (66) requiring further investigation. Third, we employed a relatively coarse brain parcellation (100 nodes) (32). Although findings were similar when using a finer 200 nodes parcellation, finer-grained analyses could increase the performance further (67). Fourth, in-scanner tasks were limited to few and not highly demanding cognitive tasks. Comparing tasks of different difficulty levels might amplify observable neural characteristics underlying intelligence (20). Fifth, our sample was restricted to 22-37 year-old subjects, leaving the question of generalizability to a broader age range for future investigations.

In summary, despite an extensive body of research predicting intelligence from brain connectivity exists, the conceptual insights provided by those studies are limited. We propose systematic functional brain connection selection as a means to increase our understanding about the neural code of individual differences in intelligence. General and crystallized intelligence were better predicted from functional connectivity than fluid intelligence, proposing differences in their underlying neural substrates. Further, prediction performance depended critically on the cognitive states during fMRI assessment with demanding tasks performing best. Notably, predictive functional brain connections were distributed across the whole brain, going beyond those proposed by major intelligence theories. Finally, we identified brain-wide networks of the most predictive 1000 connections. These depended critically on cognitive state and differed between general, crystallized, and fluid intelligence. In sum, our results suggest intelligence as emerging from global brain characteristics, rather than from isolated brain regions or single neural networks. In a broader context, our study offers a framework for future predictive modelling studies that prioritize meaningful insights into human cognition over the mere maximization of prediction performance.

## Materials and Methods

### Preregistration

Before data analysis, all analyses, sample sizes, and variables of interest were preregistered on the Open Science Framework: https://osf.io/nm7xy. Note that the study also includes not registered post-hoc analyses to further characterize brain connections identified as most relevant for intelligence prediction.

### Participants

The Human Connectome Project (HCP) Young Adult Sample S1200 including 1200 subjects of age 22–37 years (656 female, 1089 right-handed) was used for main analyses. Study procedures were approved by the Washington University Institutional Review Board, and informed consent, in accordance with the declaration of Helsinki, was obtained from all participants (31). Subjects with missing cognitive data or a mini-mental state examination score ≤ 26 (serious cognitive impairment) were excluded. Performance scores of 12 cognitive tests of the remaining 1186 subjects were used to estimate latent factors of general and fluid intelligence, and to generate composite scores of crystallized intelligence (next section). After additional exclusion due to missing fMRI data, missing in-scanner task performance scores, and excessive head motion (see below), the final sample comprised 806 subjects (418 female, 733 right-handed, 22-37 years, 28.6 years mean age).

### Intelligence

To estimate general intelligence as latent factor, bi-factor analysis (68) was performed according to Dubois et al. (19) from 12 cognitive measures (*SI Appendix*, Table S1) of 1186 subjects. Fluid intelligence was estimated as latent factor (one factor, exploratory factor analysis, oblimin rotation) from seven measures (picture sequence memory, dimensional change card sort, flanker task, Penn progressive matrices, processing speed, variable short Penn line orientation test, list sorting). Crystallized intelligence was operationalized as sum of standardized scores from the picture vocabulary and the oral reading recognition task.

### Data acquisition and preprocessing

Resting-state fMRI data (four runs) and fMRI data acquired during seven tasks (working memory, gambling, motor, language processing, relational processing, social cognition, emotion processing; two runs each) were used for analyses. Resting-state runs span 14:33 min, while task runs range from 2:16 min to 5:01 min. For general data acquisition details see Van Essen et al. (31), for information concerning the resting state refer to Smith et al. (69), and for fMRI tasks see Barch et al. (70). In brief, fMRI data were acquired with a gradient-echo EPI sequence (TR = 720 ms, TE = 33.1 ms, flip angle = 52°, 2-mm isotropic voxel resolution, multiband factor = 8) on a 3 T Siemens Skyra with a 32-channel head coil. We used the minimally preprocessed fMRI data (71). Further preprocessing steps comprised a nuisance regression strategy with 24 head motion parameters, eight mean signals from white matter and cerebrospinal fluid, as well as with four global signals (72). Global signal regression has been shown to be beneficial for removing global artifacts driven by motion and respiration, thus improving accuracies of predicting behavioral constructs (73). As task activation can produce systematic inflation of task functional connectivity estimates and task-regressed functional connectivity has been shown to more validly predict human phenotypes (74), basis-set task regressors were applied together with the other nuisance regressors to remove task-evoked neural activation (74). In-scanner head motion was measured by framewise displacement (FD) (75) and subjects were only included if mean FD < 0.2 mm, proportion of spikes (FD > 0.25 mm) < 20%, and no spikes were above 5 mm (72). Lastly, time series of neural activation were extracted from 100 regions (nodes) covering the entire cortex (32).

### Functional connectivity

Eight subject-specific weighted functional connectivity (FC) matrices (one per state: rest, seven tasks) were constructed from Fisher z-transformed Pearson correlations between the time series of neural activation from the 100 cortical regions. FC was first computed for RL and LR phase directions separately and averaged afterwards. In addition, two latent FC matrices were constructed via connection-wise factor analysis (33), i.e., one latent FC across resting state and all task states, and one latent FC across all task states. All regions were assigned to seven functional brain networks as defined in Yeo et al. (35).

### Prediction features

To compare different brain networks in their performance to predict intelligence, a functional brain connection selection approach was developed including systematic training and testing of multiple prediction models with different sets of brain connections as input features (Fig. 1*A*): a) one whole-brain model including all connections, b) seven models of within-network connections, c) 21 models of between-network connections, d) seven models with connections of all but one network, and e) seven models with connections within a specific network and between this network and all other networks. Models were trained for each state-specific FC separately, resulting in 10 (rest + 7 tasks + 2 latent) x 43 (selection-specific) models per intelligence component (general, crystallized, and fluid intelligence). Additional models were trained with different numbers (45, 190, 435, 780, and 1000 out of 4950) of randomly selected connections as well as with the same numbers of connections between randomly selected nodes (100 permutations each).

Next, we tested with five additional models the predictive performance of functional connections between clusters of brain regions proposed by prominent intelligence theories. The first model was trained on connections between eight clusters corresponding to the meta-analytically derived revised P-FIT model (13, 15). Second, two models containing connections between brain clusters specified in the Multiple-Demand (MD) theory (14) were tested: one model included connections between 13 clusters proposed by Duncan (14), while the other model comprised connections between 26 clusters from a newer version of the theory (37). Third, the role of the lateral PFC for predicting intelligence was tested with a model including functional brain connections between four clusters similar to Cole et al. (38) and a model extending these clusters to 14 lateral PFC nodes. To best capture brain clusters proposed by the revised P-FIT, the MD (Duncan), and the LPFC (Cole) theory, nodes of the Schaefer parcellation (100 nodes) (32) closest to the clusters proposed by the respective theory were selected (smallest Euclidean distance to the original coordinates). For Diachek’s MD theory, nodes best matching a mask provided by the authors and for the extended PFC theory, all lateral PFC nodes from the Schaefer parcellation were selected. The performance of these models was tested against 100 permutations of models with an equal number of connections between an equal number of nodes, which were randomly selected.

### Prediction models

For all analyses, the preregistered pipeline and parameters were used. Specifically, all prediction models were trained and tested with 5-fold cross-validation within an HCP subsample (*N*=610). Family structure was considered by grouping individuals of the same family into the same fold, while simultaneously ensuring approximately equal distributions of family-average intelligence scores via stratified folds. To control for covariates, an out-of-sample deconfounding approach was applied to avoid leakage between training-and test data. First, confounding variables were regressed out via linear regression (19) from both the target and feature variables (76) in the training sample only and residuals were z-standardized. Second, the regression coefficients and standardization parameters (*M* and *SD*) estimated in the training sample were applied to the target and feature variables of the test sample. Considered confound variables included age (Pearson correlation between age and general intelligence in the main sample: *r* = −0.13, *p* = 0.002), sex (*r* = 0.18, *p* < 0.001), handedness (*r* = 0.00, *p* = 0.998), mean framewise displacement (FD, *r* = −0.20, *p* < 0.001), and the mean number of spikes – FD > 0.2 mm (*r* = −0.20, *p* < 0.001; *SI Appendix* Table S2). Feed forward Neural Networks (implemented with Pytorch) (77) with hidden layers and Rectified Linear Unit activation functions were trained to predict intelligence scores. A learning rate of 0.01, a Mean Squared Error (MSE) loss function, a dropout of 0.25, and a Stochastic Gradient Descent (SGD) optimizer for training via backpropagation were implemented. The number of hidden layers (1–3) and the number of neurons per hidden layer (10, 50, 100) were chosen via hyperparameter optimization in a further internal cross-validation loop (3-fold). Early stopping (the stopping of the training process before completing the specified number of training epochs if a specific criterion is fulfilled) was applied to prevent overfitting (78): herein, a proportion of the training data of each fold was used as validation sample (20% of training data in 5-fold cross-validation, and 30% of training data in hyperparameter optimization). Specifically, after each training step, the performance in predicting intelligence scores of the validation sample of the respective model under training was evaluated and training was stopped if the validation loss did not decrease within the last 100 training epochs, or if the training exceeded a maximal number of training epochs (20,000).

### Interpreting prediction models with relevance back-mapping

To localize brain connections most contributing to predictions, importance of model features was evaluated with layer-wise relevance propagation (LRP) (30), a methodology ascertaining contributions of single input features to predictions based on backpropagation. During this process, shares that each model neuron has on the output are calculated back from the last layer to the input layer of the neural network (30). We applied stepwise LRP (implemented with Captum) (79) to models trained on the main sample (10 iterations with varying stratified folds). Specifically, within each fold (5-fold cross-validation), models were trained starting with all brain connections and then the most contributing connections were removed iteratively. Following, models were trained with different numbers (45 to 1000) of the most contributing connections. Finally, all models were applied to the test sample to evaluate prediction performance.

### External replication

For external replication, two independent datasets (PIOP1, PIOP2) from the AOMIC (34) were combined. Study procedures were approved by the faculty’s ethical committee (EC numbers: 2015-EXT-4366, 2017-EXT-7568), including informed consent according to the declaration of Helsinki. PIOP1 (*N*=216) contains fMRI data from six cognitive states (resting state, five tasks: emotion matching, gender-stroop, working memory, face perception, anticipation); PIOP2 (*N*=226) from four states (resting state, three tasks: emotion matching, working memory, stop signal). Replication was conducted with resting-state FC, with two task FCs (working memory, emotion matching), and two latent FCs. Again, latent FCs were estimated from resting state and all task states, and from all task states only. For latent FCs, all tasks (PIOP1 or PIOP2) were used. FMRI data were acquired with a gradient-echo EPI on a Philips 3T scanner with a 32-channel coil (3-mm isotropic voxel resolution). For face perception and resting-state scans of PIOP2, multiband scans were acquired (TR = 750 ms, TE = 28 ms, flip angle = 60°, and multiband factor = 3). For resting state of PIOP2, and for working memory, emotion matching, gender-stroop, anticipation, and stop signal scans of PIOP1 and PIOP2, sequential scans were recorded (TR = 2000 ms, TE = 28 ms, and flip angle = 76.1°). Subjects with missing imaging data, missing descriptive or behavioral data, or excessive head motion (same criteria as in the HCP) were excluded, resulting in 138 (PIOP1) and 184 (PIOP2) subjects. The minimally preprocessed fMRI data were downloaded (different preprocessing than in the HCP, fMRIprep v1.4.1) (80). Further preprocessing (extracting nuisance regressed time series) and subsequent analyses followed the same pipeline as specified previously.

### Statistics

Prediction performance was assessed using Pearson correlation (*r*) between predicted and observed intelligence scores and three error-based metrics: mean squared error (MSE), root mean squared error (RMSE), mean absolute error (MAE). For error metrics, predicted and observed intelligence scores were normalized by the range of observed scores. Statistical significance of model performance was assessed with non-parametric permutation tests and relied on Fisher z-transformed Pearson correlations between observed and predicted intelligence scores. Precisely, 100 models were trained to predict randomly shuffled intelligence scores (random assignment between subjects and scores) and the performance of these null models were evaluated against the mean performance of 10 models (varying stratified folds) trained with correct scores. Significance of model differences was assessed with non-parametric model difference permutation tests (100 permutations): differences in the prediction performance (*r*) between both models trained with the correct scores were compared to differences in performance using permuted scores. To compare the averaged prediction performance between different intelligence components and specific cognitive states, paired *t*-tests were applied (note that *t*-tests were used instead of non-parametric model difference permutation tests, as multiple models had to be compared with each other). *P*-values < 0.05 were considered as statistically significant. Note, that due to the hypotheses-generating nature of this project, we report *p*-values uncorrected for the number of prediction models in the main manuscript but for analyses across multiple networks we additionally provide results and figures with FDR-corrected significances in the Supplement (*SI Appendix* Figs. S4-S10).

Robustness and generalizability of results were assessed with a three-level validation procedure (Fig. 1*B*): first, models were trained on an HCP subsample (*N*=610) with 5-fold cross-validation (internal validation). Second, models trained on this subsample were evaluated on the lockbox sample (separated before any analyses, *N*=196, lockbox validation). Third, generalizability (external replication) was tested in two samples of the AOMIC (34). Note that because of differing fMRI tasks and the Raven’s Advanced Progressive Matrices Test (RAPM, 36 item version set II) (23) was assessed in the AOMIC samples as measure of intelligence, external replication was not possible in all specific cases.

## Data and materials availability

All data used in the current study can be accessed online under: https://www.humanconnectome.org/study/hcp-young-adult (HCP), https://doi.org/10.18112/openneuro.ds002785.v2.0.0 (AOMIC-PIOP1), and https://doi.org/10.18112/openneuro.ds002790.v2.0.0 (AOMIC-PIOP2). All analysis code used in the current study was made available by the authors: Preprocessing: https://github.com/faskowit/app-fmri-2-mat; Main analyses: https://github.com/jonasAthiele/predicting_human_cognition, https://doi.org/10.5281/zenodo.10178395.

## Acknowledgments

This work was supported by the German Research Foundation (DFG, grant HI 2185-1/1 to K.H.), by the Heinrich-Böll Foundation (funds from the Federal Ministry of Education and Research, grant P145957 to J.A.T), and by the Open Access Publication Fund of the University of Würzburg. The authors thank the Human Connectome Project, WU-Minn Consortium (Principal Investigators: David Van Essen and Kamil Ugurbil; 1U54MH091657) funded by the 16 NIH Institutes and Centers that support the NIH Blueprint for Neuroscience Research; and by the McDonnell Center for Systems Neuroscience at Washington University, for providing data of the main sample. We also thank all contributors to the Amsterdam Open MRI Collection (Principal Investigator: H. Steven Scholte) for providing data of the replication samples. Further, this research was supported in part by Lilly Endowment, Inc., through its support for the Indiana University Pervasive Technology Institute.

