## Supplementary material for "Choosing explanation over performance: Insights from machine learning-based prediction of human intelligence from brain connectivity": SI Appendix

**This PDF file includes:**

Figures S1 to S25
Tables S1 to S2
SI References

**Fig. S1.**

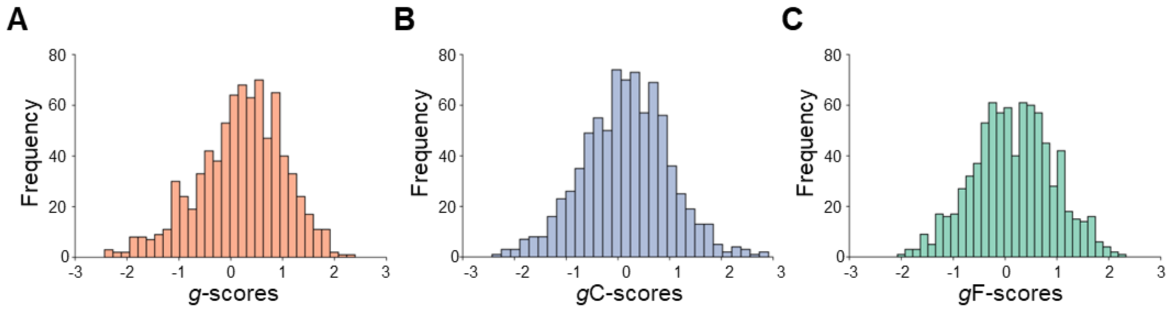


Frequency distributions of intelligence scores in the Human Connectome Project sample. Frequencies of individual intelligence scores of 806 subjects from the Human Connectome Project (main sample + lockbox sample) (1). (A) General intelligence derived as latent factor from 12 cognitive performance scores (see Table S1), (B) crystallized intelligence calculated as sum score from the performance scores of two out of 12 cognitive tasks (picture vocabulary and oral reading recognition), and (C) fluid intelligence operationalized as latent factor from the performance scores of seven out of the 12 cognitive tasks (picture sequence memory, dimensional change card sort, flanker task, Penn progressive matrices, processing speed, variable short Penn line orientation test, list sorting).

Fig. S2.

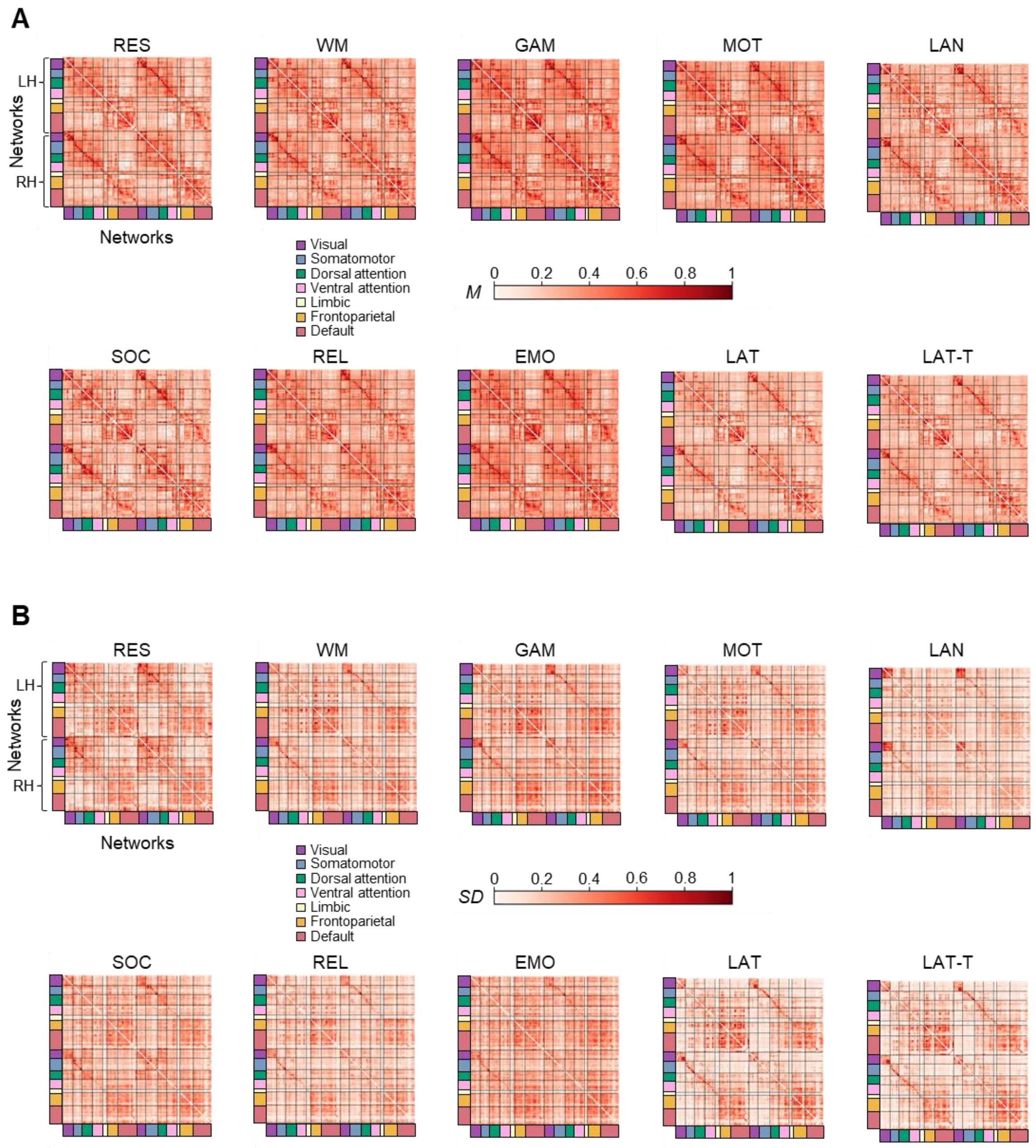


Functional connectivity in the Human Connectome Project (HCP) sample including the main sample and the lockbox sample. Means (*M*) and standard deviations (*SD*) of functional connectivity (FC) measured in 806 subjects from the HCP (1) during different cognitive states (rest, tasks, latent). Seven functional brain networks (color-coded) were defined in accordance with the Yeo atlas (2) and all 100 brain regions, as defined by the Schaefer atlas (3), were assigned to these networks. For better comparison, means and standard deviations of FC values were normalized to the range from 0 to 1. (A) Means of FC values, and (B) standard deviations of FC values. LH, left hemisphere; RH, right hemisphere; RES, resting state; WM, working memory task; GAM, gambling task; MOT, motor task; LAN, language processing task; SOC, social cognition task; REL, relational processing task; EMO, emotion processing task; LAT, latent FC of resting state and all task states; LAT-T, latent FC of all task states.

**Fig. S3.**

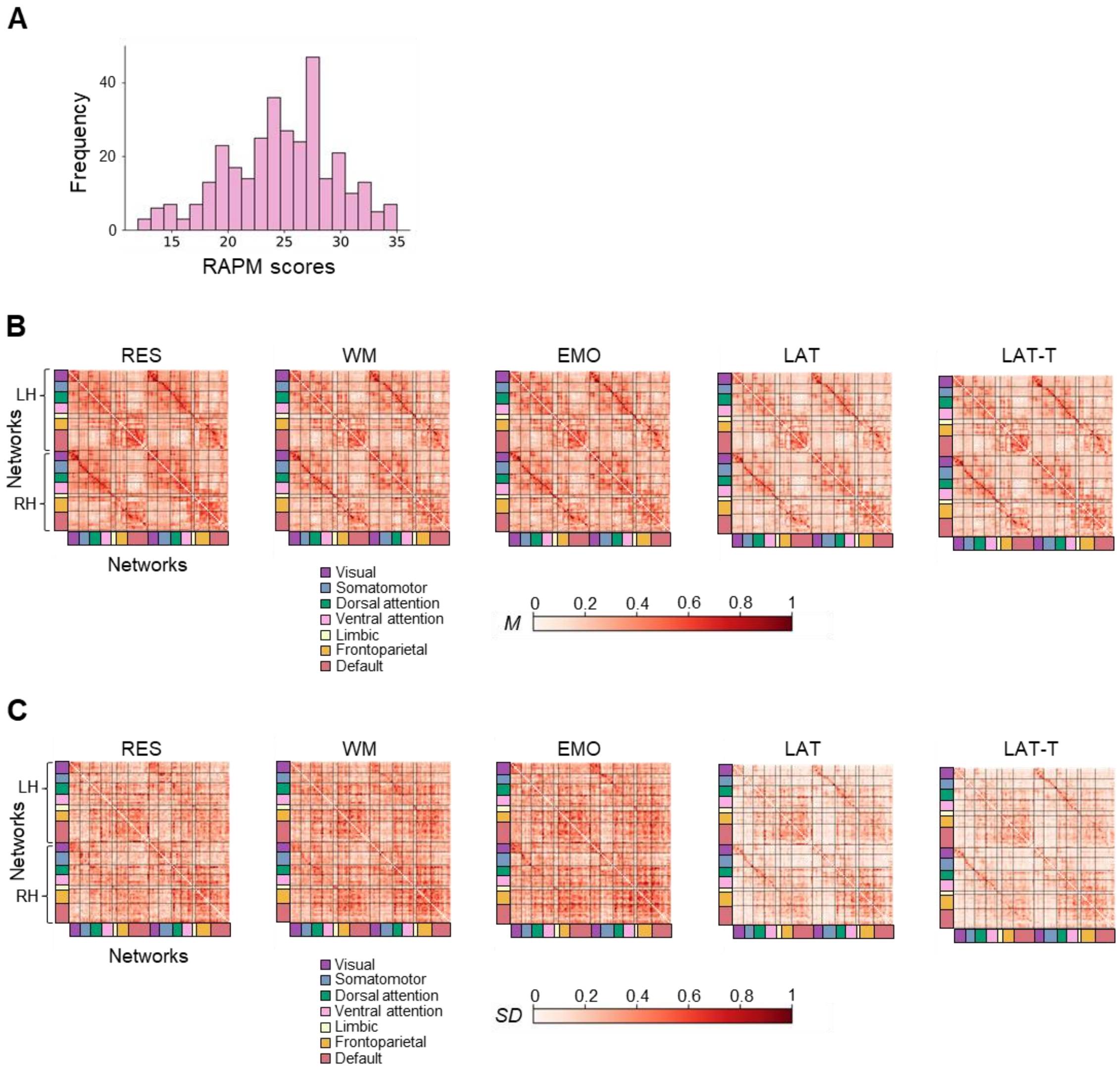


Intelligence and functional connectivity in the combined Amsterdam Open MRI Collection (AOMIC) sample (replication sample). Fluid intelligence as well as means (*M*) and standard deviations (*SD*) from functional connectivity (FC) of 322 subjects from two samples (PIOP1 and PIOP2) of the AOMIC (4). (A) Frequencies of individual intelligence scores (RAPM sum scores) (5). (B) Means, and (C) standard deviations of functional connectivity measured during different cognitive states (rest, task, latent). Seven functional brain networks (color-coded) were defined in accordance with the Yeo atlas (2) and all 100 brain regions, as defined by the Schaefer atlas (3), were assigned to these networks. For better comparison, means and standard deviations were normalized to the range from 0 to 1. LH, left hemisphere; RH, right hemisphere; RES, resting state; WM, working memory task; GAM, gambling task; MOT, motor task; LAN, language processing task; SOC, social cognition task; REL, relational processing task; EMO, emotion processing task; LAT, latent FC of resting state and all task states; LAT-T, latent FC of all task states.

**Fig. S4.**

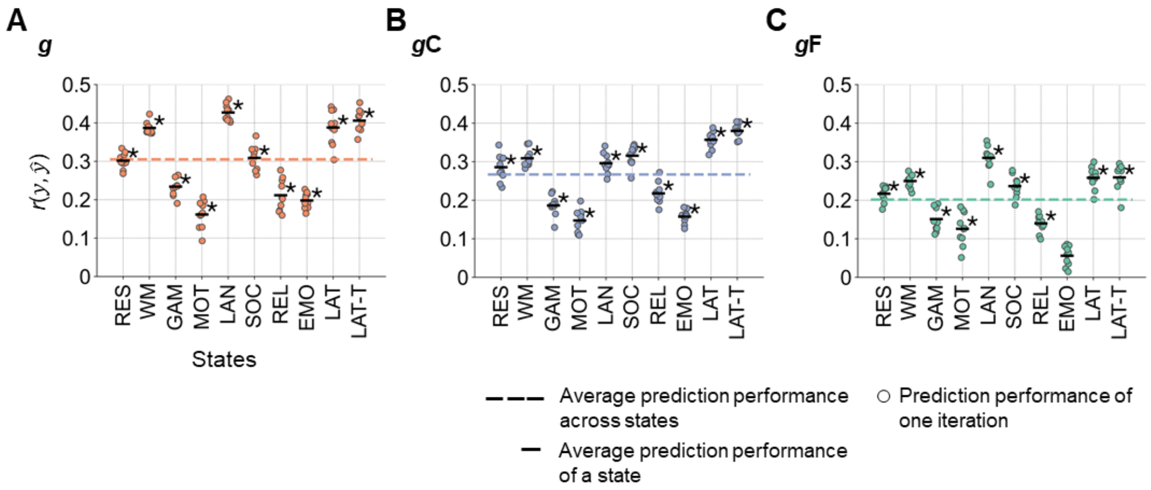


Performance of predicting intelligence from whole-brain functional connectivity in the main sample with correction for the number of prediction models. (A) Prediction of general *g*, (B) crystallized *g*C, and (C) fluid *g*F intelligence in the main sample (*N*=610, HCP). The prediction performance was calculated as Pearson correlation between observed and predicted intelligence scores $r(y, \hat{y})$. The performance of 10 prediction models trained with varying stratified folds is illustrated with colored dots (general intelligence red, crystallized intelligence blue, fluid intelligence green). The mean performance across these 10 models is indicated by the black horizontal bar. The mean performance across states is highlighted with a colored dashed line. Significant mean prediction performance ($p < 0.05$, permutation test, 100 permutations) is marked with an asterisk. *P*-values were corrected via false discovery rate (FDR) including all 10 states and three intelligence components (30 comparisons in total). RES, resting state; WM, working memory task; GAM, gambling task; MOT, motor task; LAN, language processing task; SOC, social cognition task; REL, relational processing task; EMO, emotion processing task; LAT, latent functional connectivity of resting state and all task states; LAT-T, latent functional connectivity of all task states.

**Fig. S5.**

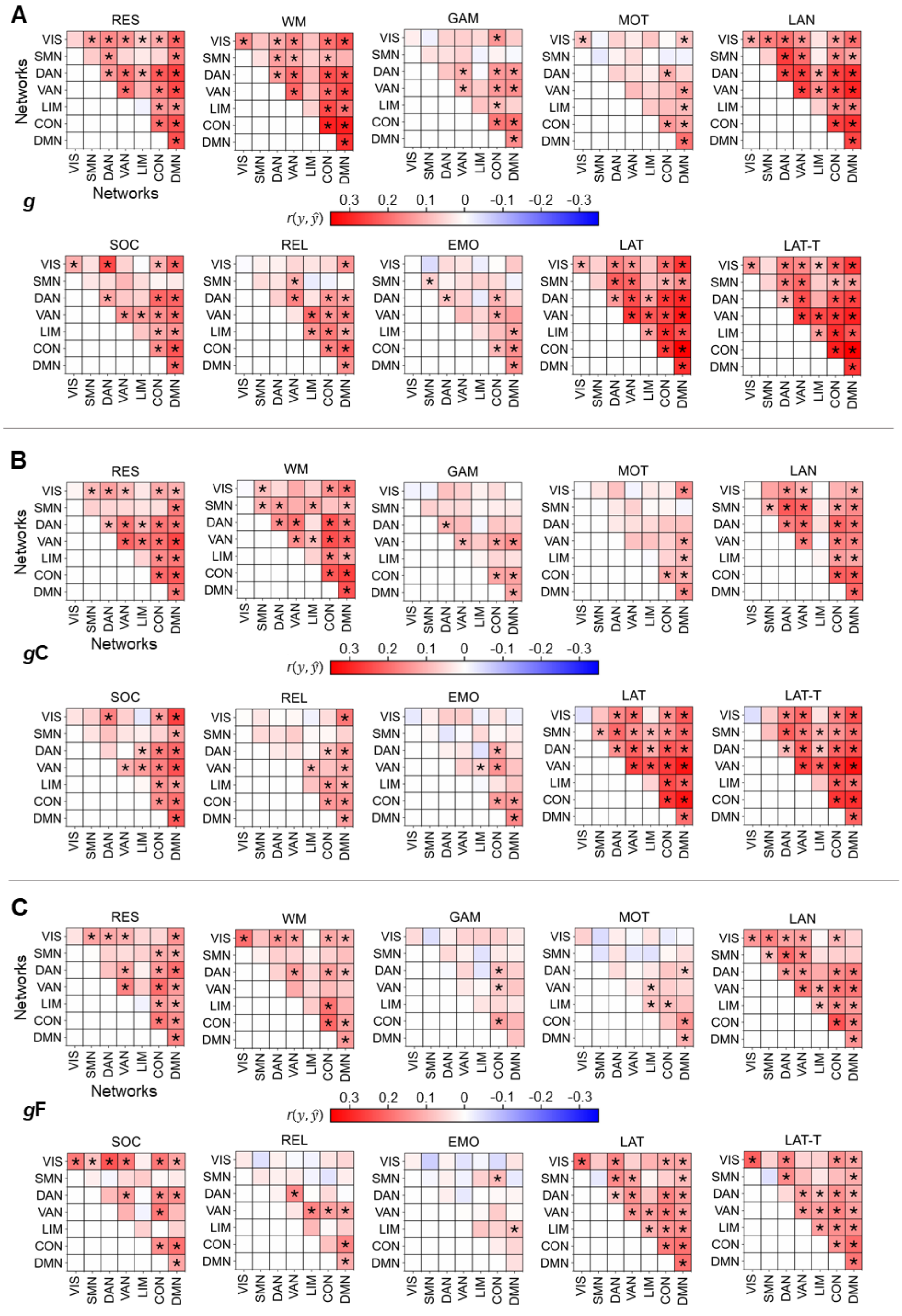


State- and network-specific performance of intelligence prediction in the main sample from functional brain connections within a specific network or between two specific networks, with correction for the number of prediction models. (A) Prediction of general *g*, (B) crystallized *g*C, and (C) fluid *g*F intelligence in the main sample (*N*=610, HCP). The prediction performance (Pearson correlation between observed and predicted intelligence scores $r(y, \hat{y})$) was calculated as average across 10 models trained with varying stratified folds. Significant prediction performance ($p < 0.05$, permutation test, 100 permutations) is marked with an asterisk. *P*-values were corrected via false discovery rate (FDR) including all 10 states, 28 network combinations, and three intelligence components (840 comparisons in total). RES, resting state; WM, working memory task; GAM, gambling task; MOT, motor task; LAN, language processing task; SOC, social cognition task; REL, relational processing task; EMO, emotion processing task; LAT, latent functional connectivity of resting state and all task states; LAT-T, latent functional connectivity of all task states; VIS, visual network; SMN, somatomotor network; DAN, dorsal attention network; VAN, salience/ventral attention network; LIM, limbic network; CON, control network; DMN, default mode network.

**Fig S6.**

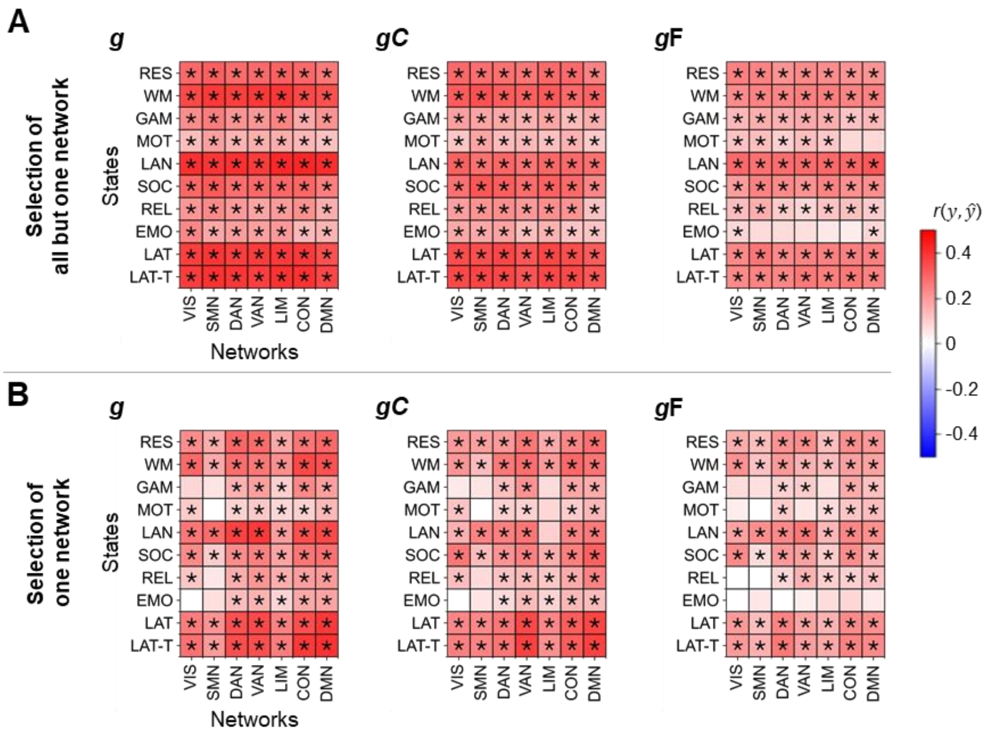


Performance of predicting intelligence in the main sample from all functional brain connections but those of one specific brain network versus from connections of one brain network only, with correction for the number of prediction models. Models for predicting general *g* (left panels), crystallized *g*C (center panels), or fluid *g*F (right panels) intelligence in the main sample (*N*=610, HCP). (A) Separate models were trained with all connections but those of one specific network. (B) Models were trained with connections of one specific network. The prediction performance (Pearson correlation between observed and predicted intelligence scores $r(y, \hat{y})$) was calculated as average across 10 models trained with varying stratified folds. Significant prediction performance ($p < 0.05$, permutation test, 100 permutations) is marked with an asterisk. *P*-values were corrected via false discovery rate (FDR) including all 10 states, 7 network combinations, and three intelligence components (210 comparisons in total). RES, resting state; WM, working memory task; GAM, gambling task; MOT, motor task; LAN, language processing task; SOC, social cognition task; REL, relational processing task; EMO, emotion processing task; LAT, latent functional connectivity of resting state and all task states; LAT-T, latent functional connectivity of all task states; VIS, visual network; SMN, somatomotor network; DAN, dorsal attention network; VAN, salience/ventral attention network; LIM, limbic network; CON, control network; DMN, default mode network.

**Fig. S7.**

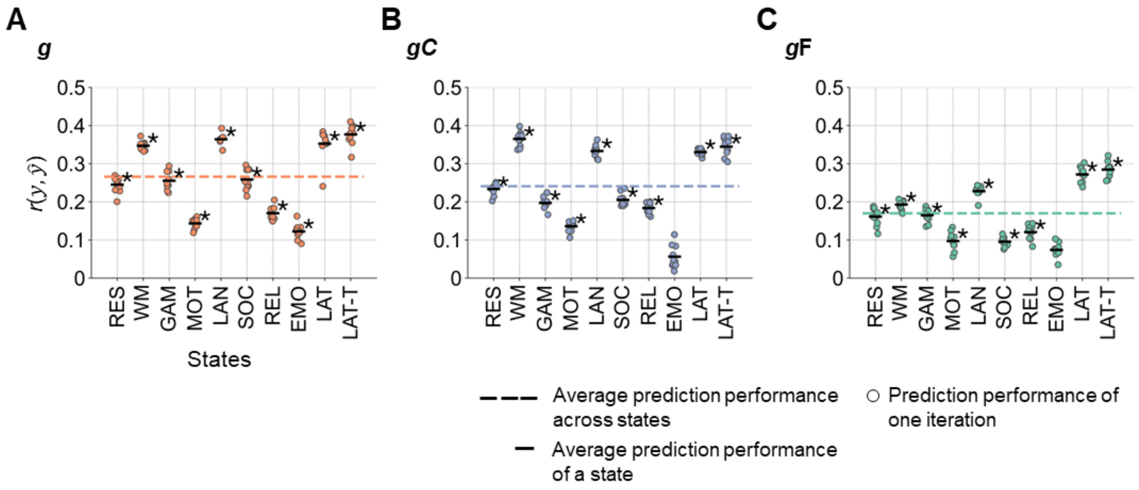


Performance of predicting intelligence in the lockbox sample with correction for the number of prediction models. Prediction models were trained on 610 subjects of the Human Connectome Project (HCP) and applied to predict intelligence scores in the lockbox sample (196 withheld subjects of the HCP). (A) Prediction of general *g*, (B) crystallized *g*C, and (C) fluid *g*F intelligence. The prediction performance was calculated as average of Pearson correlations between observed and predicted intelligence scores $r(y, \hat{y})$ across models of five folds (5-fold cross-validated training of models in the main sample) for 10 iterations with varying stratified folds. The performance of the 10 iterations is illustrated with colored dots (general intelligence red, crystallized intelligence blue, fluid intelligence green). The mean performance across models of specific states is indicated by the black horizontal bar. The mean prediction performance across states is highlighted with a colored dashed line. Significant mean prediction performance ($p <$ 0.05, tested by permutation test, 100 permutations) is marked by an asterisk. *P*-values were corrected via false discovery rate (FDR) including all 10 states and three intelligence components (30 comparisons in total). RES, resting state; WM, working memory task; GAM, gambling task; MOT, motor task; LAN, language processing task; SOC, social cognition task; REL, relational processing task; EMO, emotion processing task; LAT, latent functional connectivity of resting state and all task states; LAT-T, latent functional connectivity of all task states.

**Fig. S8.**

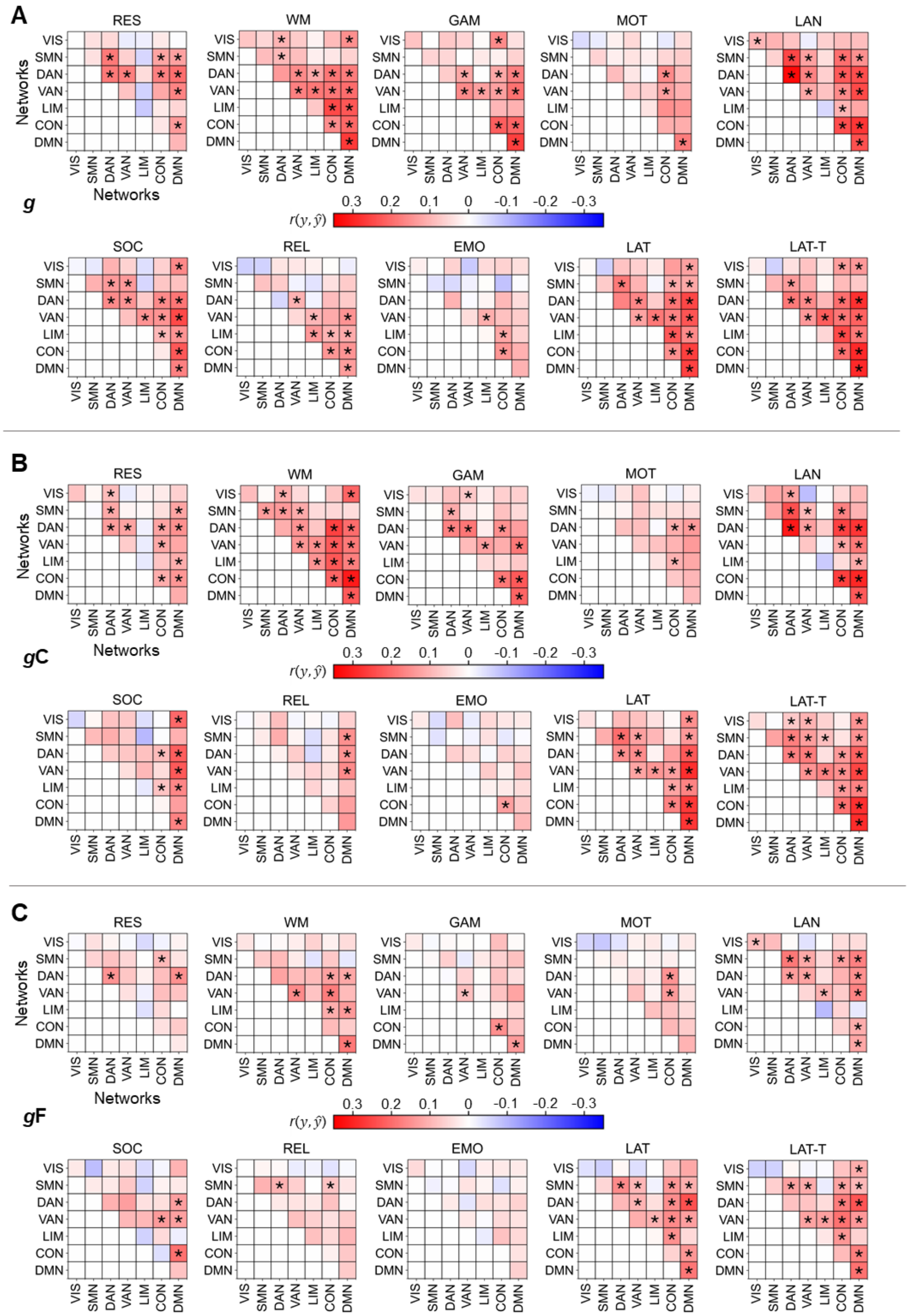


State- and network-specific performance of intelligence prediction in the lockbox sample with prediction models built in the main sample from all functional brain connections within a specific network or between two specific networks, with correction for the number of prediction models. Prediction models were trained on 610 subjects of the Human Connectome Project (HCP) and applied to predict intelligence scores of the lockbox sample (196 withheld subjects of the HCP). (A) Prediction of general *g*, (B) crystallized *g*C, and (C) fluid *g*F intelligence. The prediction performance (Pearson correlation between observed and predicted intelligence scores $r(y, \hat{y})$) was calculated as average across models of five folds (5-fold cross-validated training of models in the main sample) and 10 iterations with different stratified folds. Significant prediction performance ($p < 0.05$, tested by permutation test, 100 permutations) is marked with an asterisk. *P*-values were corrected via false discovery rate (FDR) including all 10 states, 28 network combinations, and three intelligence components (840 comparisons in total). RES, resting state; WM, working memory task; GAM, gambling task; MOT, motor task; LAN, language processing task; SOC, social cognition task; REL, relational processing task; EMO, emotion processing task; LAT, latent functional connectivity of resting state and all task states; LAT-T, latent functional connectivity of all task states; VIS, visual network; SMN, somatomotor network; DAN, dorsal attention network; VAN, salience/ventral attention network; LIM, limbic network; CON, control network; DMN, default mode network.

**Fig S9.**

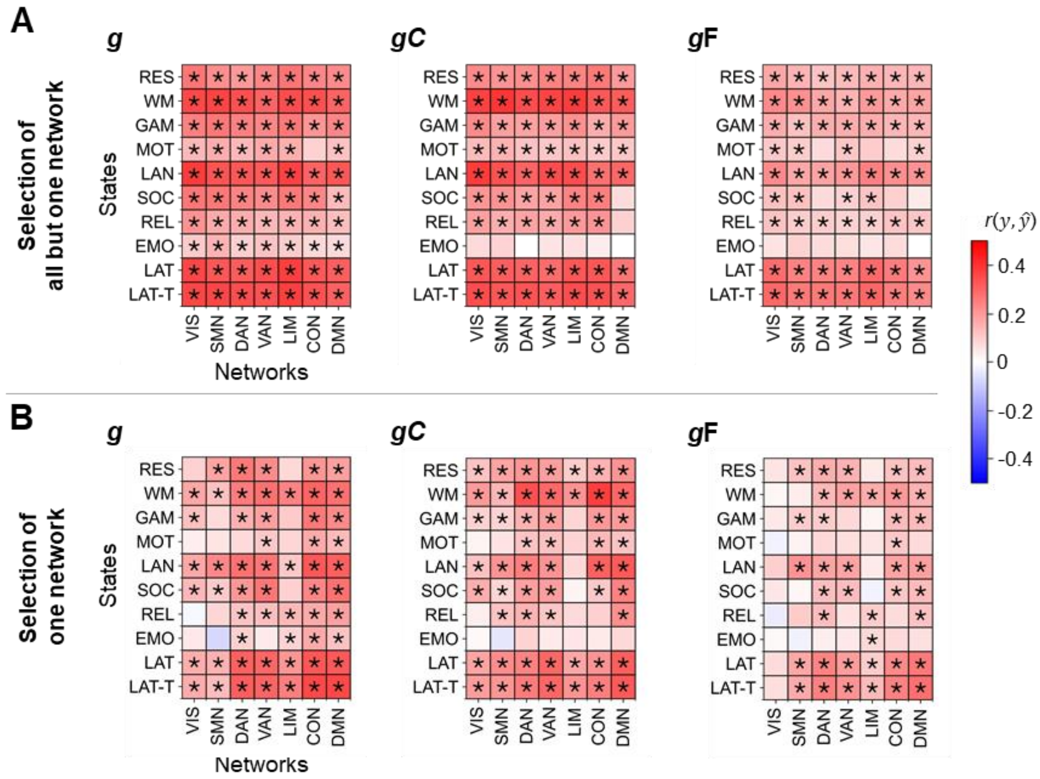


Performance of predicting intelligence in the lockbox sample with prediction models built in the main sample from all functional brain connections but those of one specific brain network versus from connections of one brain network only, with correction for the number of prediction models. Prediction models were trained on 610 subjects of the Human Connectome Project (HCP) and applied to predict intelligence scores of the lockbox sample (196 withheld subjects of the HCP). Models for predicting general *g* (left panels), crystallized *g*C (center panels), or fluid *g*F (right panels) intelligence. (A) Separate models were trained with all connections but those of one specific network. (B) Models were trained with connections of one specific network. The prediction performance (Pearson correlation between observed and predicted intelligence scores $r(y, \hat{y})$) was calculated as average across models of five folds (5-fold cross-validated training of models in the main sample) and 10 iterations with different stratified folds. Significant prediction performance ($p < 0.05$, permutation test, 100 permutations) is marked with an asterisk. *P*-values were corrected via false discovery rate (FDR) including all 10 states, 7 network combinations, and three intelligence components (210 comparisons in total). RES, resting state; WM, working memory task; GAM, gambling task; MOT, motor task; LAN, language processing task; SOC, social cognition task; REL, relational processing task; EMO, emotion processing task; LAT, latent functional connectivity of resting state and all task states; LAT-T, latent functional connectivity of all task states; VIS, visual network; SMN, somatomotor network; DAN, dorsal attention network; VAN, salience/ventral attention network; LIM, limbic network; CON, control network; DMN, default mode network.

**Fig. S10.**

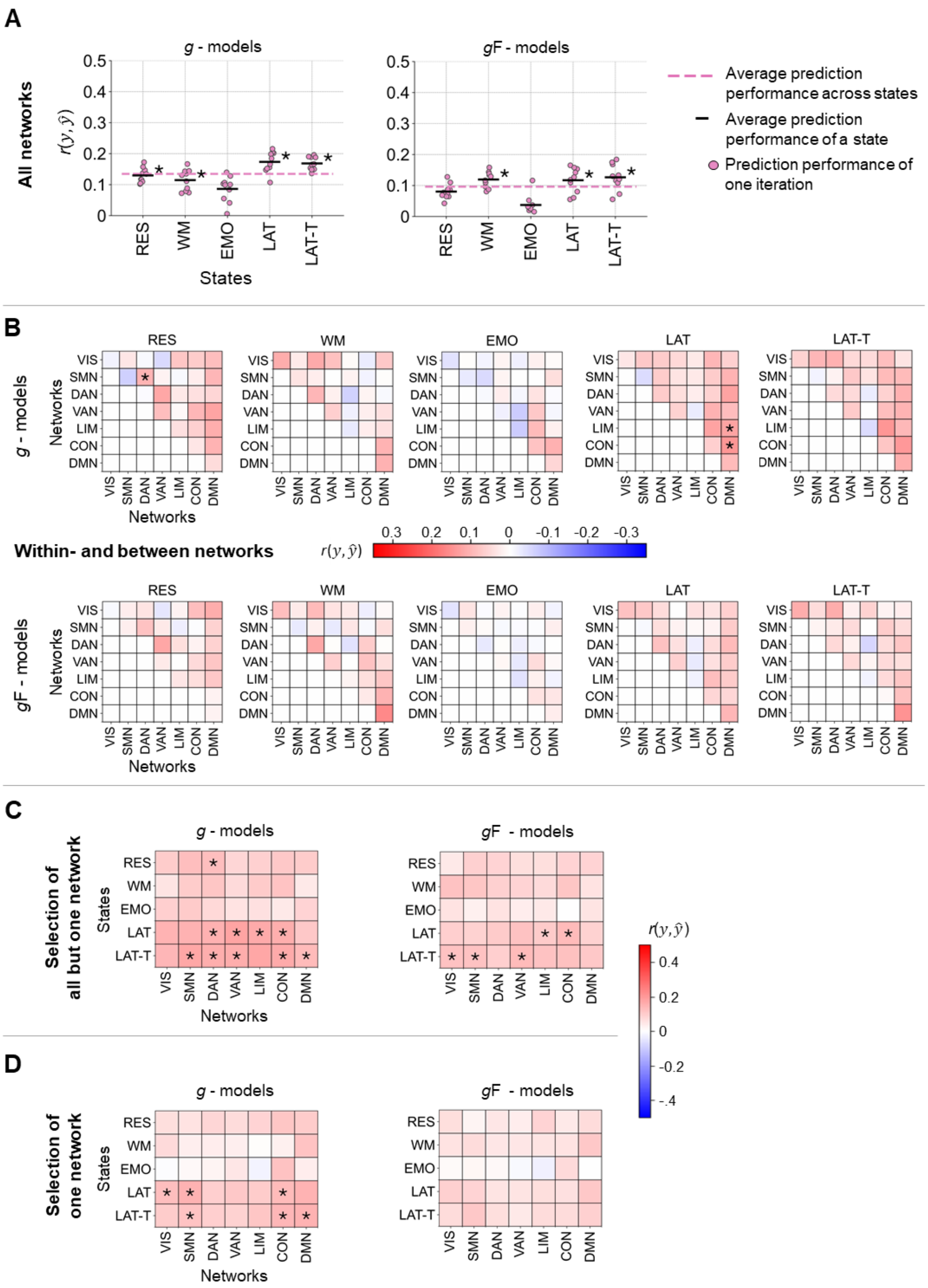


Performance of predicting intelligence in the replication sample with models trained in the main sample with correction for the number of prediction models. Models were trained on 806 subjects of the Human Connectome Project (HCP) to predict general intelligence (*g* - models) or fluid intelligence (*g*F - models) and applied to the combined samples of the Amsterdam Open MRI collection (PIOP1, PIOP2) to predict Raven's Advanced Progressive Matrices test scores (5). Separate models were trained with different selections of functional brain connections assigned to seven functional brain networks (2): (A) connections of all networks (all functional connections),

(B) connections within a specific brain network and connections between two specific brain networks, (C) connections of all networks but those of one specific network, and (D) connections of one specific network only. The prediction performance (Pearson correlation between observed and predicted intelligence scores $r(y, \hat{y})$) was calculated as average across 10 models trained with varying stratified folds. Significant prediction performance ($p < 0.05$, tested by permutation test, 100 permutations) is marked with an asterisk. *P*-values were corrected via false discovery rate (FDR) in (A) including all 5 states and two intelligence components (10 comparisons), in (B) including all 5 states, 28 network combinations, and two intelligence components (280 comparisons), and in (C) and (D) including 5 states, 7 networks, and two intelligence components (70 comparisons) respectively. RES, resting state; WM, working memory task; EMO, emotion processing task; LAT, latent functional connectivity of resting state and all task states; LAT-T, latent functional connectivity of all task states; VIS, visual network; SMN, somatomotor network; DAN, dorsal attention network; VAN, salience/ventral attention network; LIM, limbic network; CON, control network; DMN, default mode network.

**Fig. S11.**

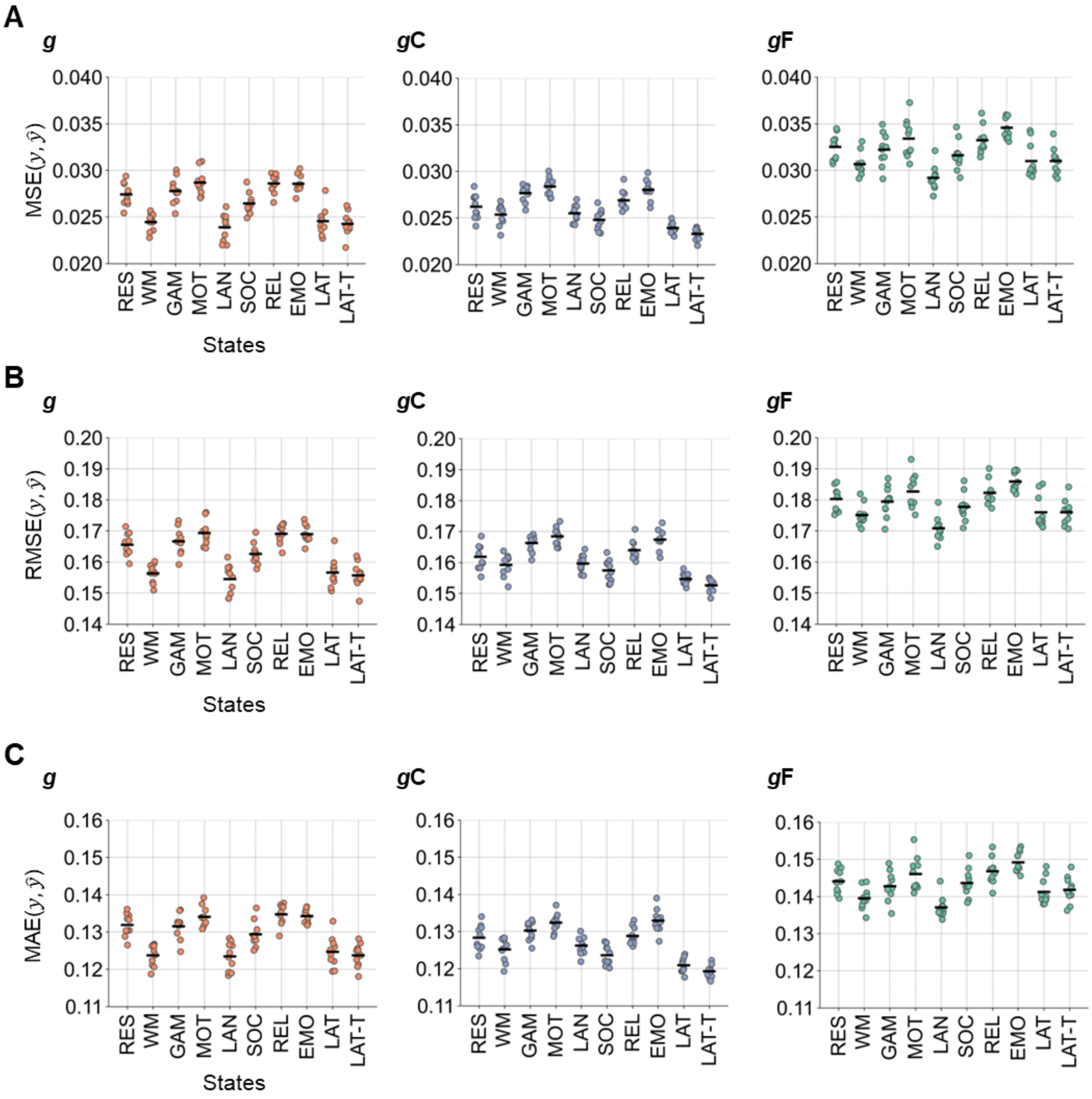


Error measures of predicting intelligence in the main sample from whole-brain functional connectivity. (A) Mean squared error (MSE), (B) root mean squared error (RMSE), and (C) mean absolute error (MAE) of predicting general *g* (left panels), crystallized *g*C (center panel), and fluid *g*F (right panel) intelligence. Error measures were calculated between observed and predicted intelligence scores from 10 iterations of model training with varying stratified sample divisions. Error measures of single prediction models are displayed with colored dots (general intelligence red, crystallized intelligence blue, fluid intelligence green). Mean error scores across the 10 models are indicated by the black horizontal bars. RES, resting state; WM, working memory task; GAM, gambling task; MOT, motor task; LAN, language processing task; SOC, social cognition task; REL, relational processing task; EMO, emotion processing task; LAT, latent functional connectivity of resting state and all task states; LAT-T, latent functional connectivity of all task states.

**Fig. S12.**

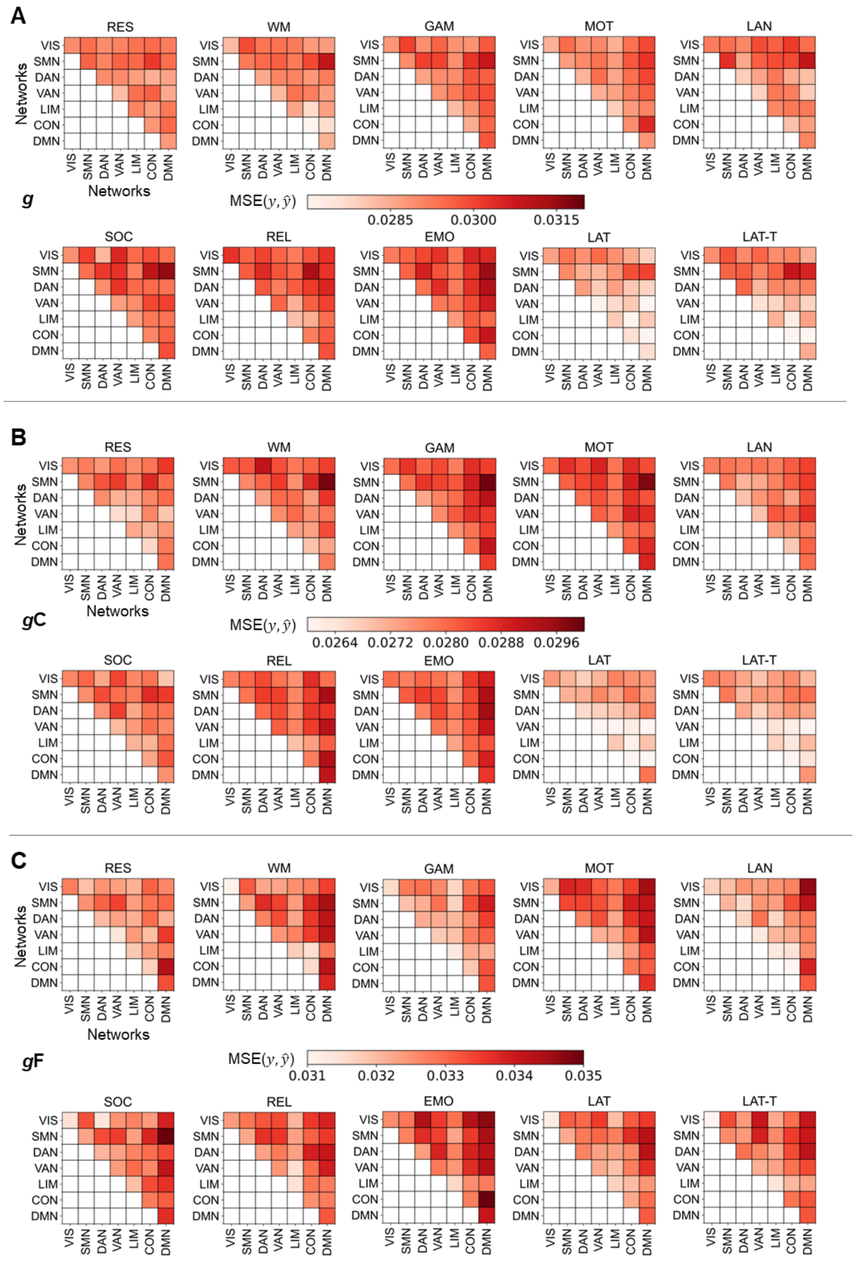

State- and network-specific mean squared errors (MSE) of intelligence prediction in the main sample from functional brain connections within a specific network or between two specific networks. (A) Prediction of general *g*, (B) crystallized *g*C, and (C) fluid *g*F intelligence. Error measures were calculated as mean squared error between observed and predicted intelligence scores $MSE(y,\hat{y})$ and averaged across 10 iterations of model training with varying stratified folds. RES, resting state; WM, working memory task; GAM, gambling task; MOT, motor task; LAN, language processing task; SOC, social cognition task; REL, relational processing task; EMO, emotion processing task; LAT, latent functional connectivity of resting state and all task states; LAT-T, latent functional connectivity of all task states; VIS, visual network; SMN, somatomotor network; DAN, dorsal attention network; VAN, salience/ventral attention network; LIM, limbic network; CON, control network; DMN, default mode network.

**Fig. S13.**

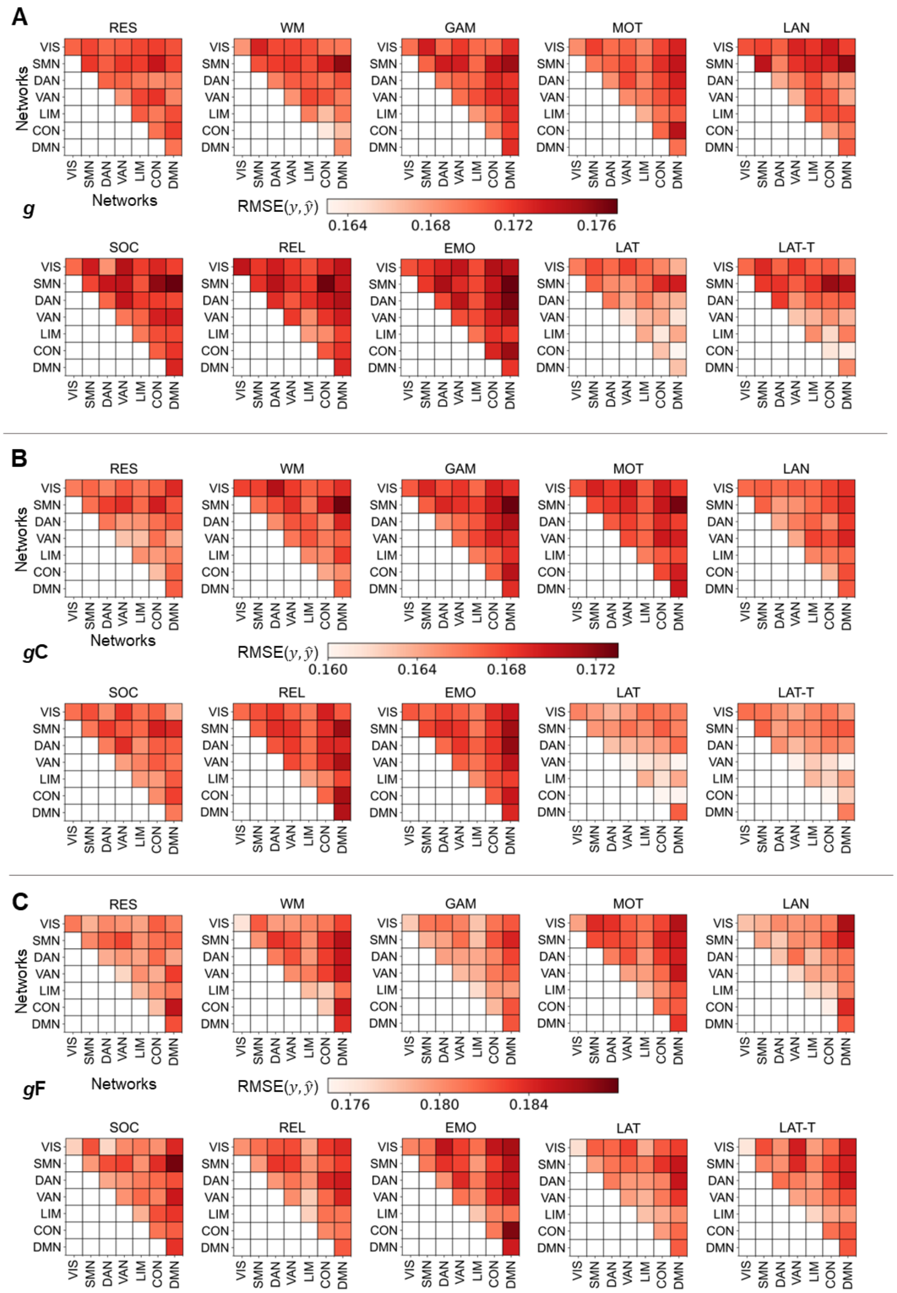

State- and network-specific root mean squared errors (RMSE) of intelligence prediction in the main sample from all functional brain connections within a specific network or between two specific networks. (A) Prediction of general *g*, (B) crystallized *g*C, and (C) fluid *g*F intelligence. Error measures were calculated as root mean squared error between observed and predicted intelligence scores $RMSE(y,\hat{y})$ and averaged across 10 iterations of model training with varying stratified folds. RES, resting state; WM, working memory task; GAM, gambling task; MOT, motor task; LAN, language processing task; SOC, social cognition task; REL, relational processing task; EMO, emotion processing task; LAT, latent functional connectivity of resting state and all task states; LAT-T, latent functional connectivity of all task states; VIS, visual network; SMN, somatomotor network; DAN, dorsal attention network; VAN, salience/ventral attention network; LIM, limbic network; CON, control network; DMN, default mode network.

**Fig. S14.**

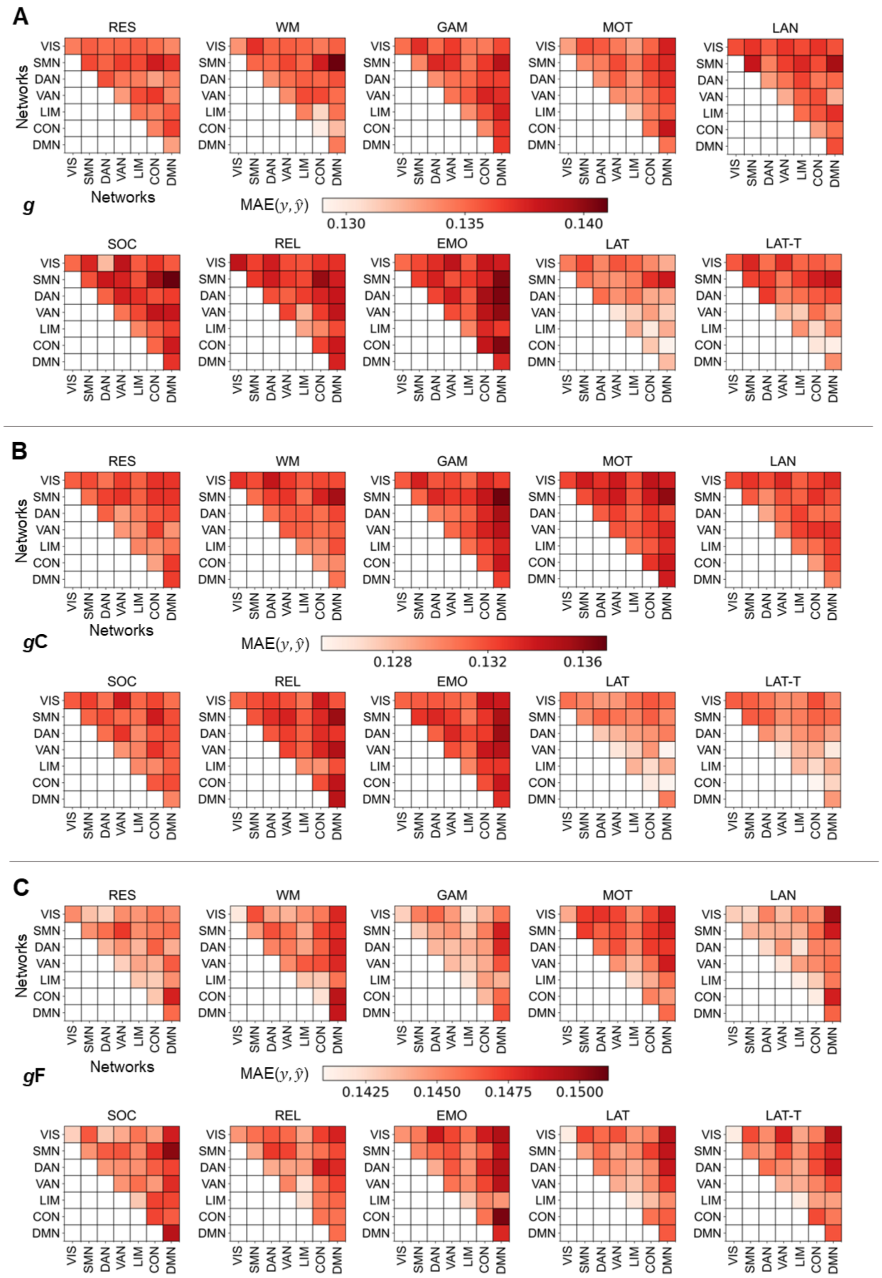


State- and network-specific mean absolute errors (MAE) of intelligence prediction in the main sample from all functional brain connections within a specific network or between two specific networks. (A) Prediction of general *g*, (B) crystallized *g*C, and (C) fluid *g*F intelligence. Error measures were calculated as mean absolute error between observed and predicted intelligence scores $MAE(y, \hat{y})$ and averaged across 10 iterations of model training with varying stratified folds. RES, resting state; WM, working memory task; GAM, gambling task; MOT, motor task; LAN, language processing task; SOC, social cognition task; REL, relational processing task; EMO, emotion processing task; LAT, latent functional connectivity of resting state and all task states; LAT-T, latent functional connectivity of all task states; VIS, visual network; SMN, somatomotor network; DAN, dorsal attention network; VAN, salience/ventral attention network; LIM, limbic network; CON, control network; DMN, default mode network.

**Fig. S15.**

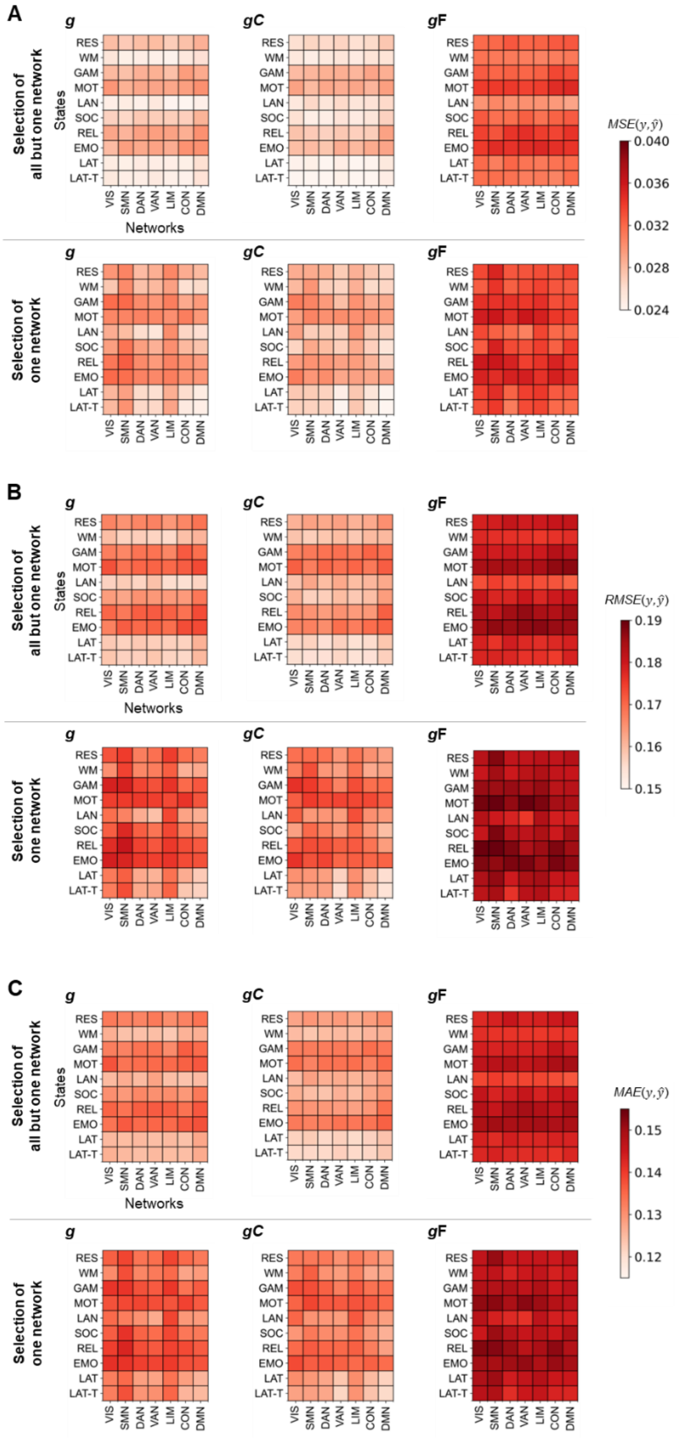


Error measures of predicting intelligence in the main sample from all functional brain connections but those of one specific brain network versus from connections of only one brain network. Models for predicting general *g* (left panels), crystallized *g*C (center panels), or fluid *g*F (right panels) intelligence. Error measures were calculated as (A) mean squared error (MSE), (B) root mean squared error (RMSE), and (C) mean absolute error (MAE) between observed $y$ and predicted $\hat{y}$ intelligence scores and averaged across 10 iterations of model training with varying stratified folds. Upper panels: Separate models were trained with all connections but those of one specific network. Lower panels: Models were trained with all connections of one specific network. RES, resting state; WM, working memory task; GAM, gambling task; MOT, motor task; LAN, language processing task; SOC, social cognition task; REL, relational processing task; EMO, emotion processing task; LAT, latent functional connectivity of resting state and all task states; LAT-T, latent functional connectivity of all task states; VIS, visual network; SMN, somatomotor network; DAN, dorsal attention network; VAN, salience/ventral attention network; LIM, limbic network; CON, control network; DMN, default mode network.

**Fig. S16.**

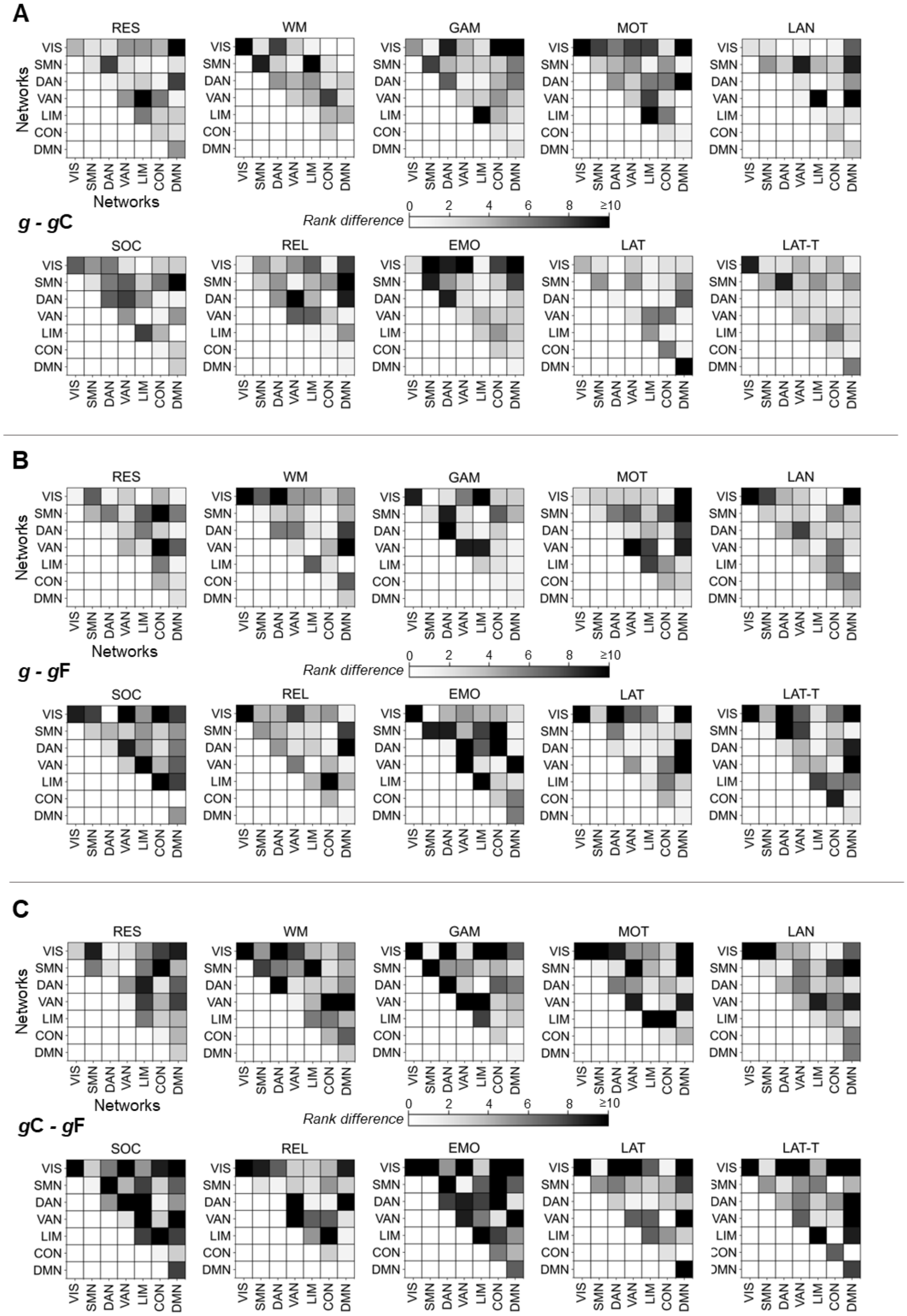


General, crystallized, and fluid intelligence vary in network-specific prediction performance in the main sample. Separate models were trained with functional brain connections within a specific brain network or between two specific networks. For each state (rest, tasks, and latent), prediction performance (see Fig. 3) was ranked from highest to lowest performance in predicting general *g*, crystallized *g*C, or fluid *g*F intelligence and the ranks were compared between the three intelligence components, resulting in rank differences for each prediction performance score of the different within- and between-network combinations. (A) Rank differences between the performance of predicting *g* and *g*C, (B) rank differences between the performance of predicting *g* and *g*F, and (C) rank differences between the performance of predicting *g*C and *g*F. Note that rank differences were computed based on the mean performance across 10 iterations of model training with varying stratified folds. RES, resting state; WM, working memory task; GAM, gambling task; MOT, motor task; LAN, language processing task; SOC, social cognition task; REL, relational processing task; EMO, emotion processing task; LAT, latent functional connectivity of resting state and all task states; LAT-T, latent functional connectivity of all task states; VIS, visual network; SMN, somatomotor network; DAN, dorsal attention network; VAN, salience/ventral attention network; LIM, limbic network; CON, control network; DMN, default mode network.

**Fig. S17.**

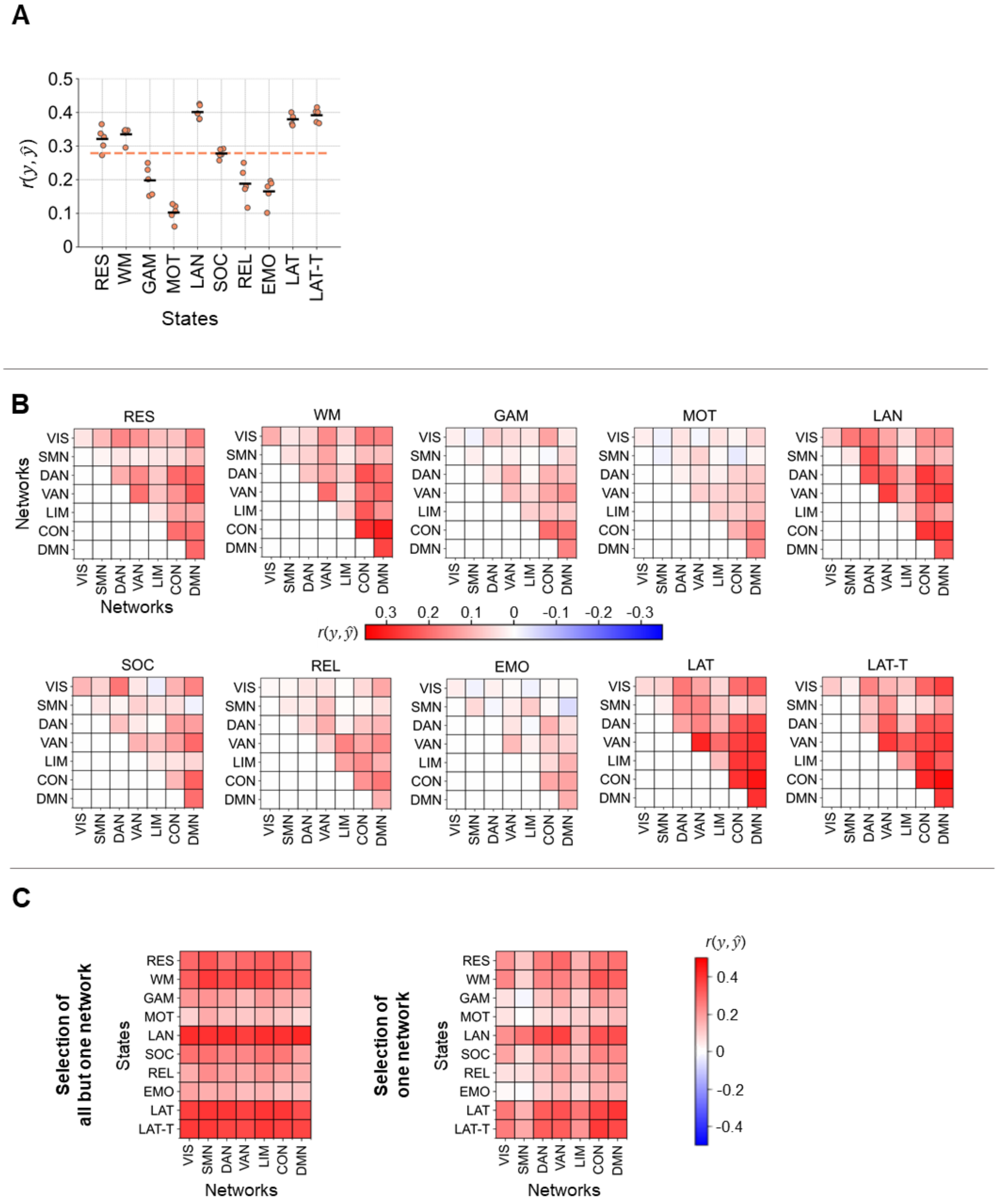


Performance of predicting intelligence in the main sample with additional control for socioeconomic status. Models were trained on 610 subjects of the Human Connectome Project (HCP) to predict general intelligence. Socioeconomic status was operationalized as household income (HCP variable: SSAGA_Income). Separate models were trained with different selections of functional brain connections assigned to seven functional brain networks: (A) connections of all networks (all functional connections), (B) connections within a specific brain network and connections between two specific brain networks, (C) connections of all networks but those of one specific network (left panel), and connections of one specific network only (right panel). The prediction performance (Pearson correlation between observed and predicted intelligence scores $r(y, \hat{y})$) was calculated as average across five models trained with varying stratified folds and is color-coded in (B) and (C). In (A), in addition to the average performance, which is here illustrated by a black horizontal bar, the performance of the five iterations is illustrated with red dots, and the mean performance across states is highlighted with a colored dashed line. Note that for this control analysis significance was not assessed. RES, resting state; WM, working memory task; EMO, emotion processing task; LAT, latent functional connectivity of resting state and all task states; LAT-T, latent functional connectivity of all task states; VIS, visual network; SMN, somatomotor network; DAN, dorsal attention network; VAN, salience/ventral attention network; LIM, limbic network; CON, control network; DMN, default mode network.

**Fig. S18.**

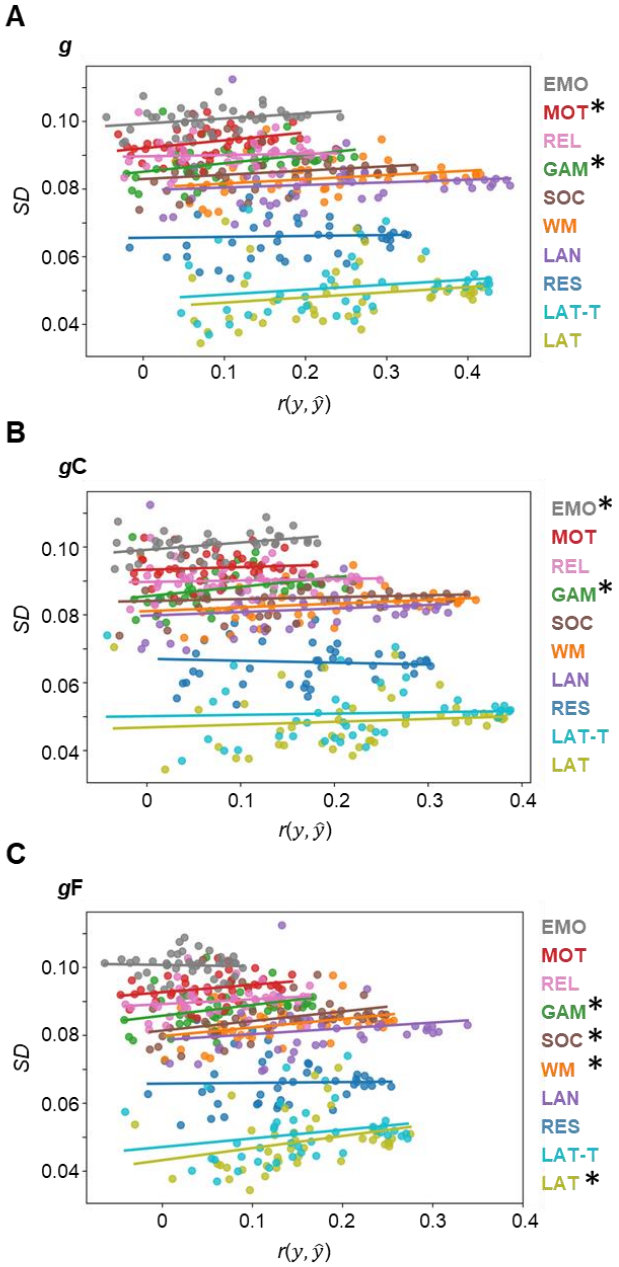


Relationship between the performance in predicting intelligence in the main sample from different brain connection selections (corresponding to different networks and network combinations) and the amount of between-subject variability within these network-specific selections of brain connections. Prediction of (A) general *g*, (B) crystallized *g*C, and C) fluid *g*F intelligence from different cognitive states. The prediction performance (x-axis) was calculated as average of Pearson correlations between observed and predicted intelligence scores $r(y, \hat{y})$ and averaged across 10 model iterations trained with varying stratified folds. The amount of variation in network-specific brain connection selections between subjects was operationalized as connection-average standard deviation (*SD*, y-axis). Brain connection selections include connections within and between seven Yeo networks (2) as well as connections of all networks but those of one specific network, and connections of one specific network only. Different cognitive states are color-coded. Significant relations ($p < 0.05$) are marked by asterisks. RES, resting state; WM, working memory task; GAM, gambling task; MOT, motor task; LAN, language processing task; SOC, social cognition task; REL, relational processing task; EMO, emotion processing task; LAT, latent functional connectivity of resting state and all task states; LAT-T, latent functional connectivity of all task states.

**Fig. S19.**

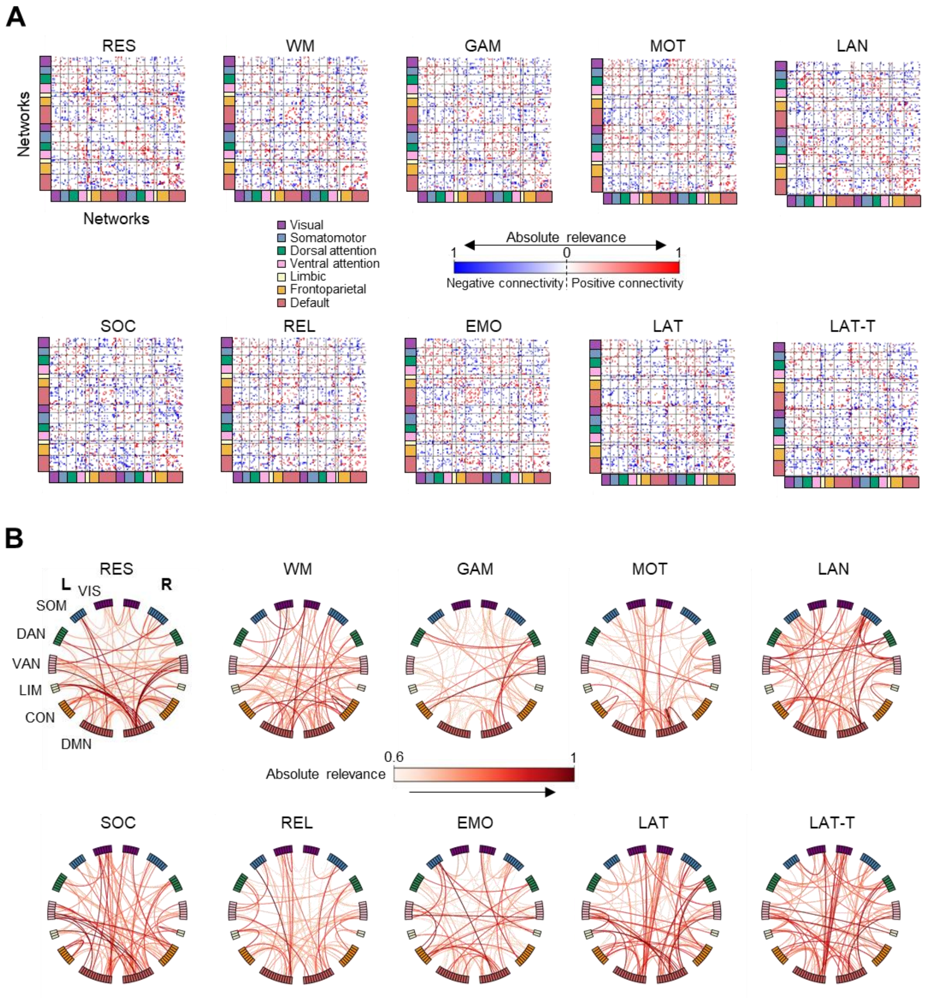


Crystallized intelligence in the main sample is best predicted from a data-driven selection of 1000 most relevant functional brain connections defining a widely distributed network. Models for predicting crystallized intelligence were trained with different numbers of the most relevant functional brain connections that were identified using stepwise layer-wise relevance propagation (LRP). (A) Matrices of the 1000 most relevant connections. Red and blue indicate whether the mean strength of a functional brain connection is positive or negative (across subjects), while the saturation of the color displays a connection's relevance in predicting crystalized intelligence (averaged across models of five folds and 10 iterations with different stratified folds). (B) Connectograms of the 100 most relevant brain connections (averaged across models of five folds and 10 iterations with different stratified folds). RES, resting state; WM, working memory task; GAM, gambling task; MOT, motor task; LAN, language processing task; SOC, social cognition task; REL, relational processing task; EMO, emotion processing task; LAT, latent functional connectivity of resting state and all task states; LAT-T, latent functional connectivity of all task states; VIS, visual network; SMN, somatomotor network; DAN, dorsal attention network; VAN, salience/ventral attention network; LIM, limbic network; CON, control/frontoparietal network; DMN, default mode network.

**Fig. S20.**

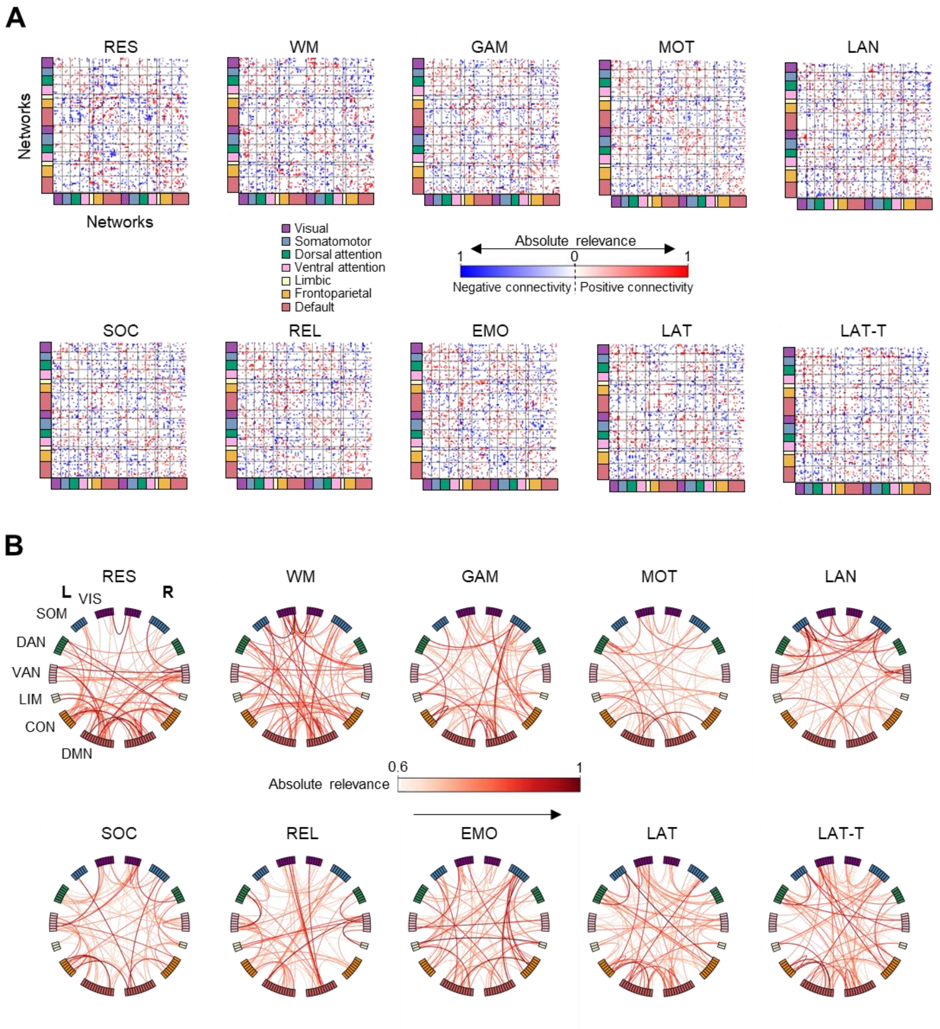


Fluid intelligence in the main sample is best predicted from a data-driven selection of 1000 most relevant functional brain connections defining a widely distributed network. Models for predicting fluid intelligence were trained with different numbers of the most relevant functional brain connections that were identified using stepwise layer-wise relevance propagation (LRP). (A) Matrices of the 1000 most relevant connections. Red and blue indicate whether the mean strength of a functional brain connection is positive or negative (across subjects), while the saturation of the color displays a connection’s relevance in predicting fluid intelligence (averaged across models of five folds and 10 iterations with different stratified folds). (B) Connectograms of 100 most relevant connections (averaged across models of five folds and 10 iterations with different stratified folds). RES, resting state; WM, working memory task; GAM, gambling task; MOT, motor task; LAN, language processing task; SOC, social cognition task; REL, relational processing task; EMO, emotion processing task; LAT, latent functional connectivity of resting state and all task states; LAT-T, latent functional connectivity of all task states; VIS, visual network; SMN, somatomotor network; DAN, dorsal attention network; VAN, salience/ventral attention network; LIM, limbic network; CON, control/frontoparietal network; DMN, default mode network.

**Fig. S21.**

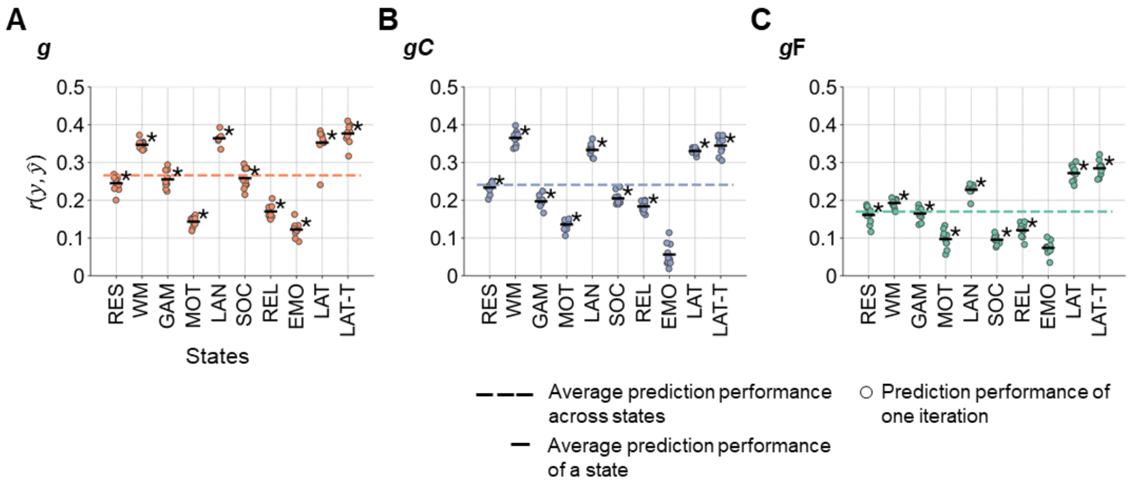


Performance of predicting intelligence in the lockbox sample with whole-brain prediction models built in the main sample. Prediction models were trained on 610 subjects of the Human Connectome Project (HCP) and applied to predict intelligence scores in the lockbox sample (196 withheld subjects of the HCP). (A) Prediction of general *g*, (B) crystallized *g*C, and (C) fluid *g*F intelligence. The prediction performance was calculated as average of Pearson correlations between observed and predicted intelligence scores $r(y, \hat{y})$ across models of five folds (5-fold cross-validated training of models in the main sample) for 10 iterations with varying stratified folds. The performance of the 10 iterations is illustrated with colored dots (general intelligence red, crystallized intelligence blue, fluid intelligence green). The mean performance across models of specific states is indicated by the black horizontal bar. The mean prediction performance across states is highlighted with a colored dashed line. Significant mean prediction performance ($p < 0.05$, tested by permutation test, 100 permutations) is marked by an asterisk (uncorrected for the number of prediction models). RES, resting state; WM, working memory task; GAM, gambling task; MOT, motor task; LAN, language processing task; SOC, social cognition task; REL, relational processing task; EMO, emotion processing task; LAT, latent functional connectivity of resting state and all task states; LAT-T, latent functional connectivity of all task states.

**Fig. S22.**

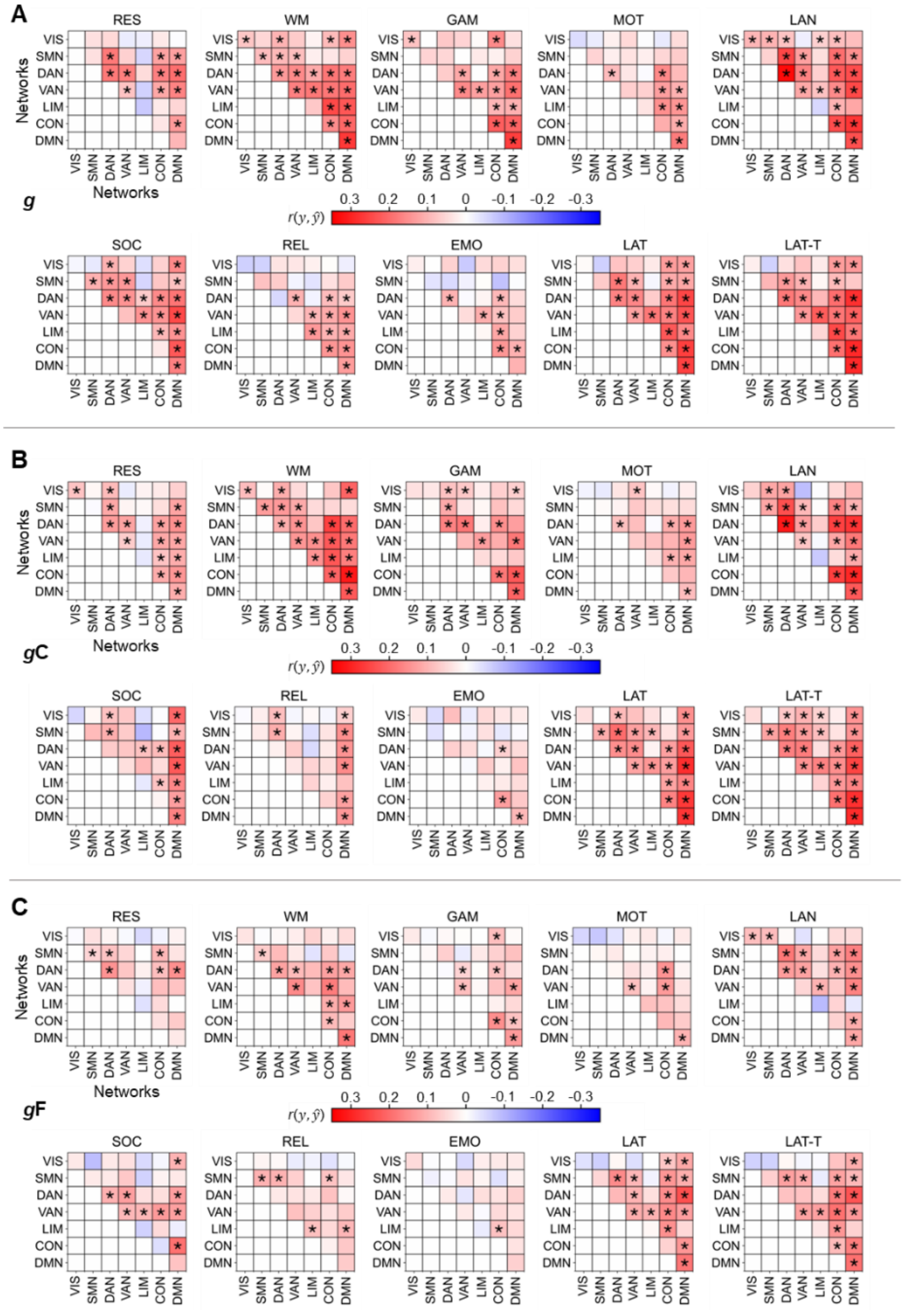

State- and network-specific performance of intelligence prediction in the lockbox sample with prediction models built in the main sample from all functional brain connections within a specific network or between two specific networks. Prediction models were trained on 610 subjects of the Human Connectome Project (HCP) and applied to predict intelligence scores of the lockbox sample (196 withheld subjects of the HCP). (A) Prediction of general *g*, (B) crystallized *g*C, and (C) fluid *g*F intelligence. Prediction performance (Pearson correlation between observed and predicted intelligence scores $r(y, \hat{y})$) was calculated as average across models of five folds (5-fold cross-validated training of models in the main sample) and 10 iterations with different stratified folds. Significant prediction performance ($p < 0.05$, tested by permutation test, 100 permutations) is marked with an asterisk (uncorrected for the number of prediction models). RES, resting state; WM, working memory task; GAM, gambling task; MOT, motor task; LAN, language processing task; SOC, social cognition task; REL, relational processing task; EMO, emotion processing task; LAT, latent functional connectivity of resting state and all task states; LAT-T, latent functional connectivity of all task states; VIS, visual network; SMN, somatomotor network; DAN, dorsal attention network; VAN, salience/ventral attention network; LIM, limbic network; CON, control network; DMN, default mode network.

**Fig. S23.**

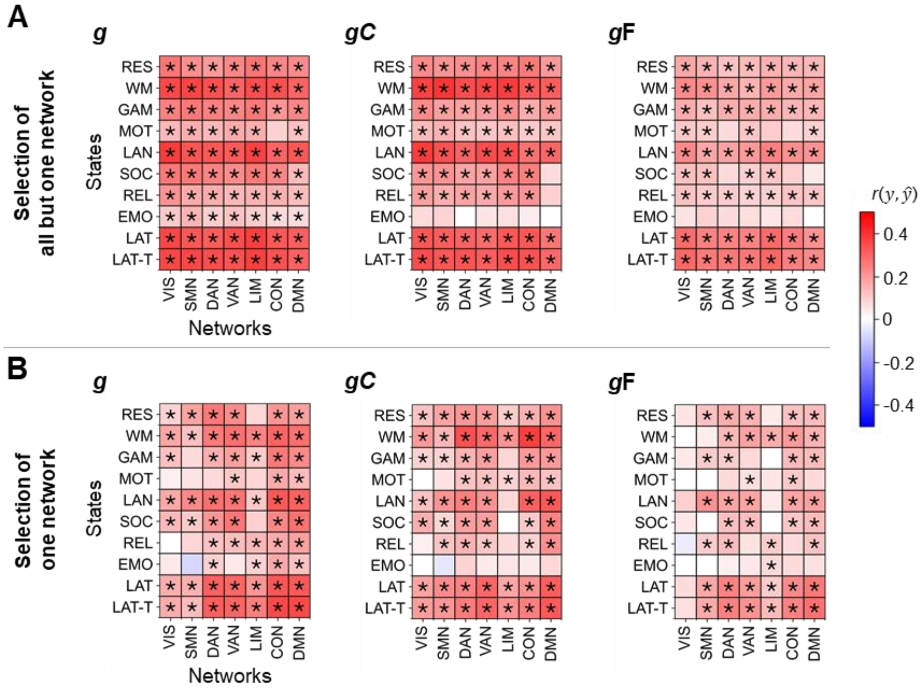


Performance of predicting intelligence in the lockbox sample with prediction models built in the main sample from all functional brain connections but those of one specific brain network versus from connections of one brain network only. Prediction models were trained on 610 subjects of the Human Connectome Project (HCP) and applied to predict intelligence scores of the lockbox sample (196 withheld subjects of the HCP). Models for predicting general *g* (left panels), crystallized *g*C (center panels), or fluid *g*F (right panels) intelligence. (A) Separate models were trained with all connections but those of one specific network. (B) Models were trained with connections of one specific network. Prediction performance (Pearson correlation between observed and predicted intelligence scores $r(y, \hat{y})$) was calculated as average across models of five folds (5-fold cross-validated training of models in the main sample) and 10 iterations with different stratified folds. Significant prediction performance ($p < 0.05$, tested by permutation test, 100 permutations) is marked with an asterisk (uncorrected for the number of prediction models). RES, resting state; WM, working memory task; GAM, gambling task; MOT, motor task; LAN, language processing task; SOC, social cognition task; REL, relational processing task; EMO, emotion processing task; LAT, latent functional connectivity of resting state and all task states; LAT-T, latent functional connectivity of all task states; VIS, visual network; SMN, somatomotor network; DAN, dorsal attention network; VAN, salience/ventral attention network; LIM, limbic network; CON, control network; DMN, default mode network.

**Fig. S24.**

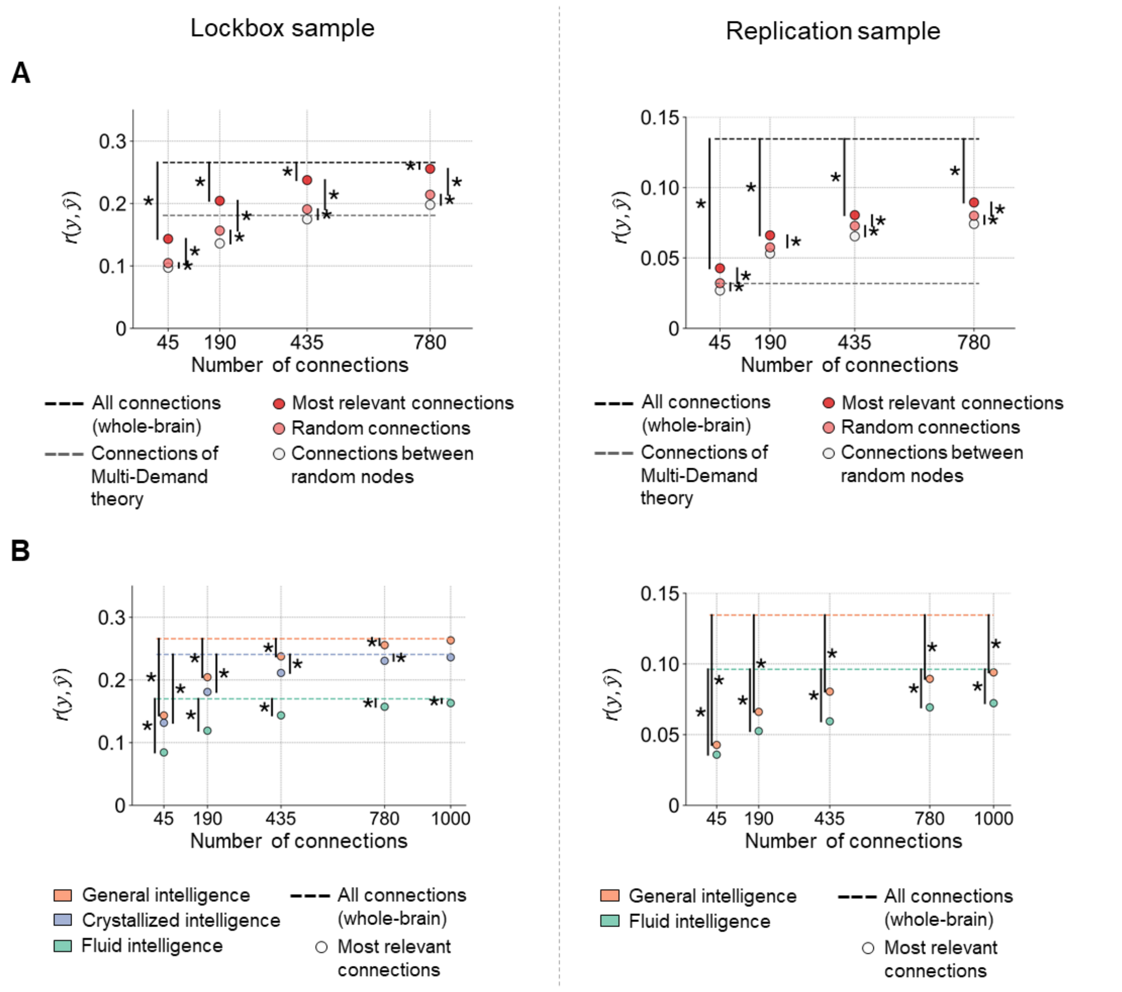


Performance of predicting intelligence in the lockbox and replication samples with prediction models build in the main sample with different selections of functional brain connections. First, models were trained with functional connectivity of subjects from the main sample of the Human Connectome Project (HCP, 610 subjects) to predict general, crystallized, or fluid intelligence scores and applied to the lockbox sample (196 withheld subjects of the HCP). Second, models trained for the prediction of general or fluid intelligence in the HCP were applied to the replication sample, i.e., combined samples of the Amsterdam Open MRI collection (PIOP1, PIOP2), to predict Raven's Advanced Progressive Matrices (RAPM) test scores (5). Prediction performance (Pearson correlation between observed and predicted intelligence scores $r(y, \hat{y})$) was calculated as average across all states (rest, tasks, latent) and across all iterations/permutations. (A) Prediction performance of main sample models trained for predicting general intelligence with different numbers of the most relevant brain connections (black circles, assessed with LRP, 10 iterations with varying stratified folds), randomly selected connections (gray circles, 100 permutations), and connections between randomly selected nodes (white circles, 100 permutations). The black dashed lines indicate the state-average performance of models trained with all functional connections (see Figs. S21 and S25*A*). The gray lines illustrate state-average performance of models trained with connections proposed by the intelligence theory that performed best in the main sample, i.e., MD by Diachek et al. (6) for reference (for clarity, significant differences to other model performance are not displayed). Significant differences ($p < 0.05$, paired $t$-test) are marked with asterisks. (B) Performance of main sample models trained with different numbers of the most relevant functional brain connections for predicting general (red dots), crystallized (blue dots), or fluid (green dots) intelligence scores. The most relevant functional brain connections were identified using stepwise LRP during model training in the main sample. Dashed lines illustrate the state-average prediction performance of models trained with all functional connections (see Figs. S21 and S25*A*). Significant differences ($p < 0.05$, paired *t*-test) between the prediction performance of specific numbers of most relevant connections and whole-brain predictions are marked with asterisks.

**Fig. S25.**

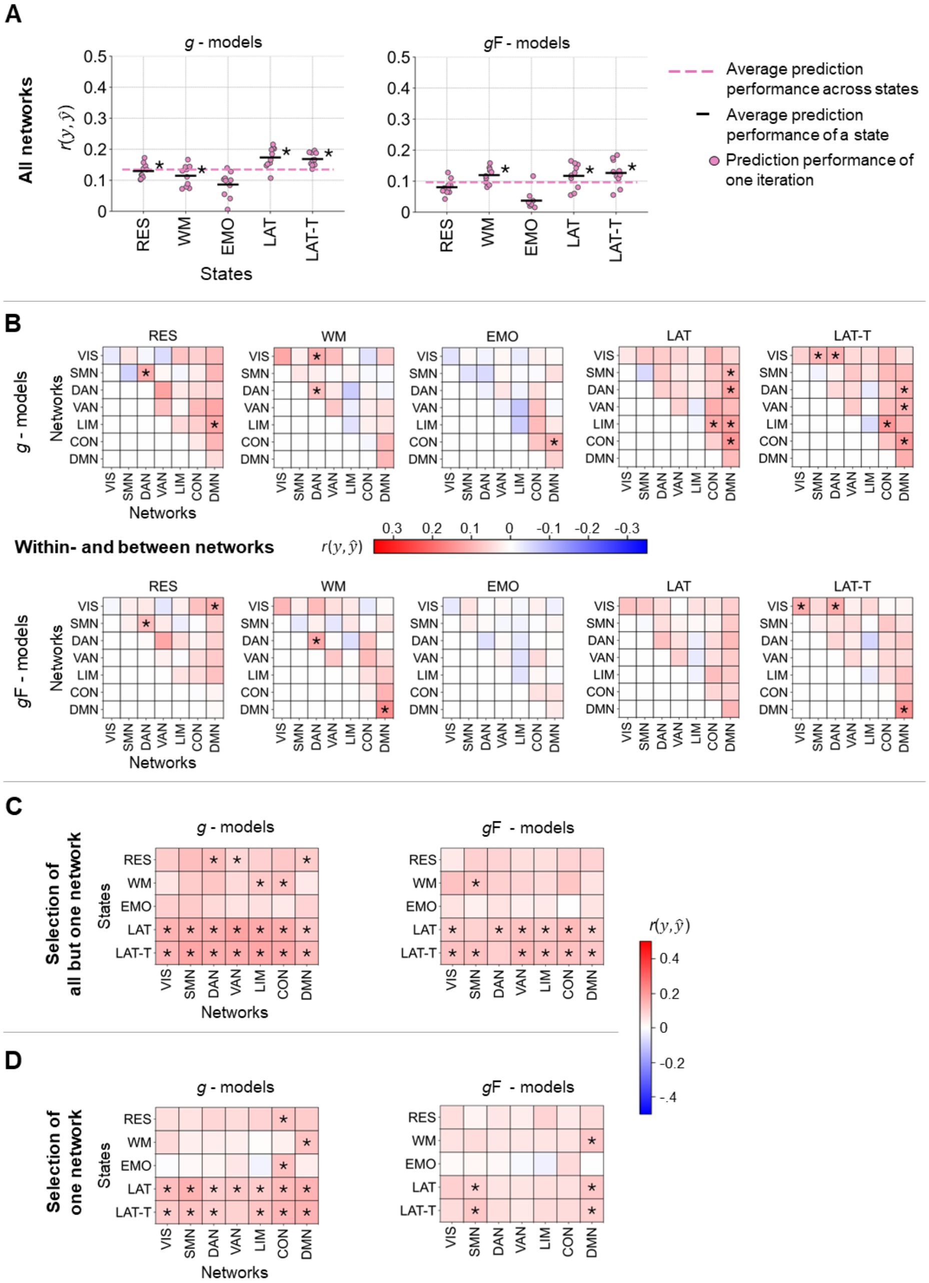


Performance of predicting intelligence in the replication sample with models trained in the Human Connectome Project sample. Models were trained on 806 subjects of the Human Connectome Project (HCP) to predict general intelligence (*g* - models) or fluid intelligence (*g*F - models) and applied to the combined samples of the Amsterdam Open MRI collection (PIOP1, PIOP2) to predict Raven's Advanced Progressive Matrices test scores (5). Separate models were trained with different selections of functional brain connections assigned to seven functional brain networks (2): (A) connections of all networks (all functional connections), (B) connections within a specific brain network and connections between two specific brain networks, (C) connections of all networks but those of one specific network, and (D) connections of one specific network only. Prediction performance (Pearson correlation between observed and predicted intelligence scores

$r(y, \hat{y})$) was calculated as average across 10 models trained with varying stratified folds. Significant prediction performance ($p < 0.05$, tested by permutation test, 100 permutations) is marked with an asterisk (uncorrected for the number of prediction models). RES, resting state; WM, working memory task; EMO, emotion processing task; LAT, latent functional connectivity of resting state and all task states; LAT-T, latent functional connectivity of all task states; VIS, visual network; SMN, somatomotor network; DAN, dorsal attention network; VAN, salience/ventral attention network; LIM, limbic network; CON, control network; DMN, default mode network.

**Table S1.** Cognitive measures from the Human Connectome Project (HCP) (1) used to estimate a latent factor of general intelligence (*g*) and fluid intelligence (*g*F) as well as a composite score of crystallized intelligence (*g*C).

| **Test** | **Instrument** | **Measure used** |
|---|---|---|
| 1 | Episodic Memory (Picture Sequence Memory) | PicSeq_Unadj |
| 2 | Executive Function/Cognitive Flexibility (Dimensional Change Card Sort) | CardSort_Unadj |
| 3 | Executive Function/Inhibition (Flanker Task) | Flanker_Unadj |
| 4 | Fluid Intelligence (Penn Progressive Matrices) | PMAT24_A_CR |
| 5 | Language/Reading Decoding (Oral Reading Recognition) | ReadEng_Unadj |
| 6 | Language/Vocabulary Comprehension (Picture Vocabulary) | PicVocab_Unadj |
| 7 | Processing Speed (Pattern Completion Processing Speed) | ProcSpeed_Unadj |
| 8 | Self-Regulation/Impulsivity (Delay Discounting) | DDisc_AUC_200 + DDisc_AUC_40K |
| 9 | Spatial Orientation (Variable Short Penn Line Orientation Test) | VSPLOT_TC |
| 10 | Sustained Attention (Short Penn Continuous Performance Test) | $\frac{SCPT\_TP + SCPT\_TN}{(SCPT\_TP + SCPT\_TN + SCPT\_FP + SCPT\_FN)SCPT\_TPRT}$ |
| 11 | Verbal Episodic Memory (Penn Word Memory Test) | IWRD_TOT |
| 12 | Working Memory (List Sorting) | ListSort_Unadj |

**Table S2.** Pearson correlation between intelligence components and covariates in the main sample, lockbox sample and replication sample.

| **Main sample (HCP, *N* = 610)** | | | | | |
|---|---|---|---|---|---|
| | Age | Sex | Handedness | Mean framewise-displacement | Mean number of spikes |
| *g* | -0.13 (0.002) | 0.18 (< 0.001) | 0.00 (0.998) | -0.20 (< 0.001) | -0.20 (< 0.001) |
| *g*F | -0.22 (< 0.001) | 0.13 (0.001) | -0.01 (0.871) | -0.22 (< 0.001) | -0.19 (< 0.001) |
| *g*C | 0.03 (0.400) | 0.11 (0.006) | 0.02 (0.623) | -0.13 (0.001) | -0.15 (< 0.001) |
| **Lockbox sample (HCP, *N* = 196)** | | | | | |
| | Age | Sex | Handedness | Mean framewise-displacement | Mean number of spikes |
| *g* | -0.06 (0.384) | 0.21 (0.003) | 0.00 (0.998) | -0.20 (0.005) | -0.11 (0.134) |
| *g*F | -0.20 (0.006) | 0.18 (0.013) | -0.01 (0.892) | -0.16 (0.029) | -0.10 (0.156) |
| *g*C | 0.06 (0.409) | 0.17 (0.020) | 0.03 (0.715) | -0.18 (0.010) | -0.12 (0.089) |
| **Replication sample (AOMIC, *N* = 322)** | | | | | |
| | Age | Sex | Handedness | Mean framewise-displacement | Mean number of spikes |
| RAPM sum score | 0.11 (0.041) | 0.04 (0.460) | 0.03 (0.568) | -0.05 (0.420) | -0.07 (0.218) |

For each sample (main, lockbox, replication sample), the Pearson correlation *r* between the intelligence components of the respective sample (general intelligence *g*, fluid intelligence *g*F, and crystallized intelligence *g*C, Raven's Advanced Progressive Matrices RAPM test scores) and each confounding variable (age, sex, handedness, mean framewise displacement, mean number of spikes) is listed together with the *p*-value in the format *r* (*p*).
